# Mendelian Randomization Analysis Dissects the Relationship between NAFLD, T2D, and Obesity and Provides Implications to Precision Medicine

**DOI:** 10.1101/657734

**Authors:** Zhipeng Liu, Yang Zhang, Sarah Graham, Roger Pique-Regi, Xiaocheng Charlie Dong, Y. Eugene Chen, Cristen Willer, Wanqing Liu

## Abstract

**Background:** Non-alcoholic fatty liver disease (NAFLD) is epidemiologically correlated with both type 2 diabetes (T2D) and obesity. However, the causal inter-relationships among the three diseases have not been completely investigated.

**Aim:** We aim to explore the causal relationships among the three diseases.

**Design and methods:** We performed a genome-wide association study (GWAS) on fatty liver disease in ∼400,000 UK BioBank samples. Using this data as well as the largest-to-date publicly available summary-level GWAS data, we performed a two-sample bidirectional Mendelian Randomization (MR) analysis. This analysis tested the causal inter-relationship between NAFLD, T2D, and obesity, as well as the association between genetically driven NAFLD (with two well-established SNPs at the PNPLA3 and TM6SF2 loci) and glycemic and lipidemic traits, respectively. Transgenic mice expressing the human PNPLA3 I148I (TghPNPLA3-I148I) and PNPLA3 I148M (TghPNPLA3-I148M) isoforms were used to further validate the causal effects.

**Results:** We found that genetically instrumented hepatic steatosis significantly increased the risk for T2D (OR=1.3, 95% CI: [1.2, 1.4], *p*=8.3e-14) but not the intermediate glycemic phenotypes at the Bonferroni-adjusted level of significance (*p*<0.002). There was a moderate, but significant causal association between genetically driven hepatic steatosis and decreased risk for BMI (β=- 0.027 SD, 95%CI: [−0.043, −0.01], *p*=1.3e-4), but an increased risk for WHRadjBMI (Waist-Hip Ratio adjusted for BMI) (β=0.039 SD, 95%CI: [0.023, 0.054], *p*=8.2e-7), as well as a decreased level for total cholesterol (β=-0.084 SD, 95%CI [−0.13, −0.036], *p*=6.8e-4), but not triglycerides (β=0.02 SD, 95%CI [−0.023, 0.062], *p*=0.36). The reverse MR analyses suggested that genetically driven T2D (OR=1.1, 95% CI: [1.0, 1.2], *p*=1.7e-3), BMI (OR=2.3, 95% CI: [2.0, 2.7], *p*=1.4e-25) and WHRadjBMI (OR=1.5, 95% CI: [1.3, 1.8], *p*=1.1e-6) causally increase the NAFLD risk. In the animal study, as compared to the TghPNPLA3-I148I controls, the TghPNPLA3-I148M mice developed higher fasting glucose level and reduced glucose clearance. Meanwhile, the TghPNPLA3-I148M mice demonstrated a reduced body weight, increased central to peripheral fat ratio, decreased circulating total cholesterol as compared to the TghPNPLA3-I148I controls.

**Conclusion:** This large-scale bidirectional MR study suggests that lifelong, genetically driven NAFLD is a causal risk factor for T2D (hence potentially a “NAFLD-driven T2D” subtype) and central obesity (or “NAFLD-driven obesity” subtype), but protects against overall obesity; while genetically driven T2D, obesity, and central obesity also causally increase the risk of NAFLD, hence a “metabolic NAFLD”. This causal relationship revealed new insights into disease subtypes and provided novel hypotheses for precision treatment or prevention for the three diseases.

## INTRODUCTION

Non-alcoholic Fatty Liver Disease (NAFLD) is characterized by the presence of excess hepatic fat accumulation (≥ 5%) without significant alcohol use, hepatitis virus infection, or other secondary causes of hepatic fat accumulation^1^. The spectrum of NAFLD ranges from simple non-alcoholic fatty liver (NAFL) to non-alcoholic steatohepatitis (NASH), which over time can lead to cirrhosis, hepatocellular carcinoma, and organ failure ^2^. Compelling observational epidemiological studies have shown that NAFLD is highly correlated with metabolic disorders such as type 2 diabetes (T2D) ^3-5^ and obesity ^1,6,7^. All three diseases together affect over 50% of the U.S. population ^8-10^.

Dissecting the causal relationship between the three diseases is crucial for both understanding the disease etiology and developing effective therapeutic or preventive strategies. However, observational associations are limited in elucidating the causality due to various confounding factors (e.g. lifestyle, socioeconomic status) or reverse causation bias ^11^. As a result, the three diseases are often treated as comorbidities for each other in various biomedical research settings. On the other hand, current prevention and treatment strategies focus on each of the three diseases individually. Without clearly knowing the causality among the three diseases, treatments of preventive interventions may often lead to conflicting research findings and inconsistent responses to disease prevention and treatment among patients.

Mendelian randomization (MR) analysis, which uses genetic variants as proxies for the risk factors of interest, has been widely applied in understanding the causal relationship between various risks factors and human diseases, e.g. estimation of the causal effect of plasma HDL cholesterol on myocardial infarction risk ^12^. Since during the process of meiosis the alleles of the parents are randomly segregated to the offspring, the MR method is considered to be analogous to a randomized controlled trial (RCT) but less likely to be influenced by confounding factors and reverse causation ^13^. Bidirectional MR is an extension of traditional MR in which the exposure-outcome causal relationship was explored from both sides. The bidirectional framework provides an efficient way to ascertain the direction of a causal relationship, which helps alleviate the potential bias from reverse causation ^14^.

Recent MR studies have partially explored the causal relationships among the three diseases. Dongiovanni et al. showed genetically instrumented hepatic steatosis was associated with insulin resistance and a small increase in T2D risk ^15^. A study by De Silva et al. indicated that genetically raised circulating ALT and AST increased the risk of T2D^16^. Stender et al. found that the genetic predictors of BMI were associated with NAFLD^17^.However, a systematic bidirectional MR study leveraging the latest GWAS data is particularly needed to elucidate the causal relationships among the three metabolic diseases under a uniformed setting. In addition, experimental analysis e.g. animal models with a characterized natural history under controlled conditions would also help further establish the causality.

In this study, we first aimed to explore whether NAFLD casually increases risks for T2D, obesity and their related intermediate traits. We then investigated the reverse relationships, i.e. whether T2D and obesity causally affect NAFLD risk. Further, we constructed a transgenic mice model expressing human PNPLA3 isoforms, a known genetic NAFLD model to test the causal effects of hepatic perturbations on T2D and obesity.

## METHODS AND MATERIALS

### Ethics statement

The summary-level GWAS data used for mendelian randomization analyses are publicly available ^18-27^ (**Figure 1 and Supplemental Table 1**). Therefore, no specific ethical approval is required. The study of the transgenic mice experiments has been reviewed and approved by the IACUC of the Indiana University School of Medicine. This research has been conducted using the UK Biobank Resource under application number 24460.

**Figure 1.**
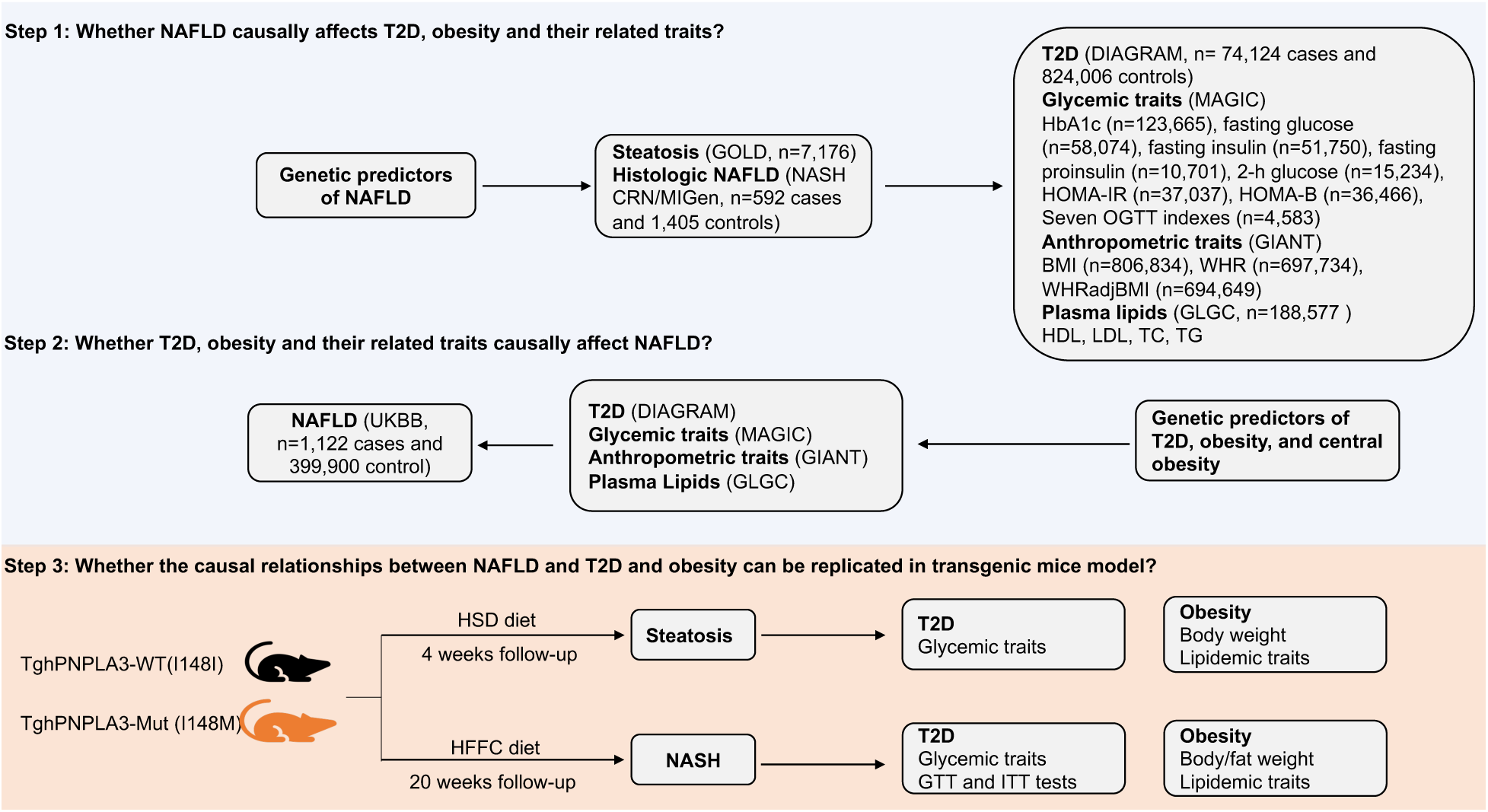
Flowchart of the study design. The summary-level associations were taken from the following genomics consortium: GOLD (Genetics of Obesity-related Liver Disease) for computerized tomography (CT) measured hepatic steatosis ^18^; NASH Clinical Research Network (NASH CRN) and Myocardial Infarction Genetics Consortium (MIGen) for biopsy-proven NAFLD ^18^; DIAbetes Genetics Replication And Meta-analysis (DIAGRAM) for T2D ^19^; Meta-Analyses of Glucose and Insulin-related traits (MAGIC) consortium for glycemic traits including HbA1c ^20^, fasting glucose ^21^, fasting insulin ^21^, fasting proinsulin ^22^, 2-h glucose ^23^, homeostatic model assessment of insulin resistance (HOMA-IR) ^24^ and β-cell function (HOMA-B) ^24^, and seven insulin secretion and action indices during oral glucose tolerance test (OGTT) ^25^; The Genetic Investigation of ANthropometric Traits (GIANT) consortium for body mass index (BMI), waist-hip ratio (WHR), and WHR adjusted for BMI (WHRadjBMI) ^26^; The Global Lipids Genetics Consortium (GLGC) for plasma high-density lipoprotein cholesterol (HDL), low-density lipoprotein cholesterol (LDL), total cholesterol (TC), and triglycerides (TG) levels ^27^.

### MR analyses

#### GWAS summary data

The summary statistics of association with computerized tomography (CT) measured hepatic steatosis were taken from a meta-analysis of 7,176 individuals by the Genetics of Obesity-related Liver Disease (GOLD) consortium ^18^. The results of the association with histologic NAFLD were taken from the same study, in which 592 biopsy-proven NAFLD patients from NASH Clinical Research Network (NASH CRN) and 1,405 controls from Myocardial Infarction Genetics Consortium (MIGen) were involved. The full GWAS summary data of this study are not publicly available, therefore only the results of the top GWAS loci associated with steatosis and histologic NAFLD were extracted. The full GWAS summary statistics of NAFLD were generated in UK Biobank (UKBB) samples consisting of 1,122 cases and 399,900 controls. The details of the GWAS of NAFLD in UKBB are described in the section below.

We downloaded full GWAS summary data of 22 glycemic and obesity traits from the largest published studies as of March 2019. These traits include T2D, glycated hemoglobin A1c (HbA1c), fasting glucose, fasting insulin, fasting proinsulin, 2-h glucose, homeostatic model assessment of insulin resistance (HOMA-IR), β-cell function (HOMA-B) and seven insulin secretion and action indices during oral glucose tolerance test (OGTT) including area under the curve of insulin levels (AUCins), ratio of AUC insulin and AUC glucose (AUCins/AUCgluc), incremental insulin at 30 min (Incre30), insulin response to glucose during the first 30 min adjusted for BMI (Ins30adjBMI), insulin sensitivity index (ISI), corrected insulin response (CIRadjISI), disposition index (DI), body mass index (BMI), waist-hip ratio (WHR), WHR adjusted for BMI (WHRadjBMI), and four plasma lipid levels. Only association results from participants of European descent were used in the present study. The details on the phenotype information, sample size, and PubMed ID of the original study are summarized in **Supplemental Table 1**.

#### GWAS of NAFLD and T2D in UK Biobank

NAFLD was defined based on ICD-9 571.8 “Other chronic nonalcoholic liver disease”) and ICD-10 K76.0 [“Fatty (change of) liver, not elsewhere classified”] from inpatient hospital diagnosis within the UK Biobank dataset. Individuals with Hepatitis B or C or with other known liver diseases (e.g. liver transplant, hepatomegaly, jaundice, or abnormal liver function study results) were excluded from the analysis. The white British subset of UK Biobank was used for analysis in SAIGE ^28^ with sex, birth year, and 4 principle components as covariates.

#### Genetic predictors selection

We used the two strongest genetic predictors of NAFLD, Patatin-like phospholipase domain-containing protein 3 (*PNPLA3*) rs738409 and Transmembrane 6 superfamily member 2 (*TM6SF2*) rs58542926, as the proxy for hepatic steatosis and histologic NAFLD. These two variants have been consistently shown to be associated with the whole spectrum of the NAFLD ^18,29-33^. Multiple MR analyses have previously been performed using these two variants to test the causal relationship between NAFLD and diseases such as Vitamin D deficiency ^34^, ischaemic heart disease ^35^, and liver damage ^15^. As *TM6SF2* rs58542926 is not genotyped in most of the GWAS summary data used in this study, rs2228603 at the *NCAN* gene locus, which is in strong linkage disequilibrium with *TM6SF2* rs58542926 (pairwise R^2^ = 0.76 based on the Phase 3 data of the 1000 Genomes Project in European individuals) and significantly associated with both hepatic fat content and NAFLD histology ^36^, was used as the proxy for *TM6SF2* rs58542926.

For the 22 glycemic and obesity traits, we selected the significant and independent genetic predictors in two steps: we first obtained all the variants that passed the genome-wide association significance level of *p*<5e-8. Then the independent genetic predictors were identified by clumping the top GWAS loci through PLINK 1.9 (https://www.cog-genomics.org/plink2) with the threshold of R^2 < 0.1 in a 500-kb window. The linkage disequilibrium was estimated based on the European samples in phase 3 of the 1000 Genomes Project ^37^. We omitted 10 traits including 2h glucose, HOMA-IR, HOMA-B, and seven OGTT traits due to lack of enough significant and independent genetic variants (number of valid variants < 3). Therefore, 12 traits were analyzed for the causal effects on NAFLD.

The instrument strength of each genetic predictor was assessed by 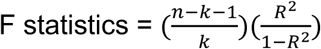, where n represents sample size, k represents the number of genetic variants, and R^2^ is the proportion of phenotypic variance explained by the genetic variants. The F statistics of all the genetic predictors in the present study were larger than the empirical threshold of 10 ^38^ (**Table 2** and **Supplemental Table 2**).

#### MR estimation

We set the exposure increasing allele as the effect allele. If the effect size was reported on the alternative allele in the original study, we multiplied the reported effect size by −1 for harmonization of the effect allele. For palindromic SNPs, we checked the reported allele frequency to avoid potential strand flipping issues ^39^. Palindromic SNPs with minor allele frequency larger than 0.42 or lack of allele frequency information were considered as ambiguous and thus removed for MR estimation. We only kept SNPs that had association results in both exposure and outcome GWAS studies.

For MR estimation with NAFLD as the exposure, we used the inverse variance weighted (IVW) method to estimate the combined causal effect of *PNPLA3* and *TM6SF2 (NCAN)* variants by assuming a fixed-effect model ^40^. As a sensitivity analysis, Wald’s method ^41^ was used to estimate the causal effect with each of the genetic variant respectively. We considered a significant causal relationship if the directions of the estimates by *PNPLA3, TM6SF2 (NCAN)*, and the two variants combined were consistent, and the combined estimate passed the Bonferroni-adjusted significance of *p*<0.05.

For the MR estimation with NAFLD as the outcome, besides the IVW method, we estimated the causal effects using additional methods including weighted median estimator ^42^ and MR-Egger ^43^ as a sensitivity analysis. IVW method provides greater precision in MR estimates in the absence of directional pleiotropy ^44^. Weighted median estimator relaxes the assumption by requiring at least 50% of the genetic variants to be valid instruments. MR-Egger provides unbiased MR estimates even if the genetic variants exhibit pleiotropic effects given the impendence between instrument strength and pleiotropic effects.

To assess the heterogeneity and identify horizontal pleiotropic outliers, we used the Q’ statistics ^45^ with modified second order weights and MR-PRESSO global test ^46^. If the horizontal pleiotropy is significant, MR-PRESSO was used to identify outliers at *p*<0.05. We then removed the outliers and retested if the causal relationship and pleiotropic effects were significant. We considered as significant if the directions of the estimates by three methods (IVW, weighted median, and MR-Egger) were consistent, IVW method passed the Bonferroni-adjusted significance of *p*<0.05, and no significant pleiotropy tested by MR-PRESSO global test and modified Q’ statistics. MR analyses were performed with “MendelianRandomization” ^47^, “MRPRESSO” ^46^, and “RadialMR” ^48^ packages in R version 3.5.0 (http://www.r-project.org/).

#### Sample overlap

Participant overlap in two-sample MR might lead to inflated Type 1 error rates due to the weak instrument bias ^49^. We examined the samples used to estimate the genetic correlations with exposure and outcome, respectively. The maximum sample overlapping rate was then calculated as the percentage of overlapped samples in the larger dataset ^49^. For example, if sample size of study 1 is n1, sample size of study 2 is n2, sample size of the overlapped cohort in study 1 is m1, sample size of the overlapped cohort in study 2 is m2, then the maximum overlapping rate = min(m1,m2) / max(n1, n2). As shown in **Supplemental Table 3**, the degree of overlap in most MR analyses is very low (<1%). For the analyses with substantial overlap (>1%), we have used robust instruments (F statistics ranges from 53 to 110) for MR estimate. Based on a simulation study ^49^, overlapping bias is unlikely to affect the results given instruments of this degree of strength.

### Animal experiments

#### Construction of transgenic mice and induction of the NAFLD phenotypes

The human PNPLA3-I148I or PNPLA3-I148M transgenic mice (TghPNPLA3-I148I or TghPNPLA3-I148M) were generated by using the PiggyBac transgenic technology at Cyagen (USA). The human BAC clone CTD-2316P10 were modified to generate the PNPLA3-I148I and PNPLA3148M isoforms and transduced into the embryo stem cell of the C57BL/6 mouse, respectively, without knocking out the mPNPLA3. All animal experiments were carried out at the animal facility with the approval from the Institutional Animal Care and Use Committee of Indiana University School of Medicine in accordance with National Institutes of Health guidelines for the care and use of laboratory animals. Six- to eight-week-old male mice were allowed for free access to water and fed regular chow (Teklad Diets 2018SX: 24% calories from protein, 18% calories from fat, and 58% calories from carbohydrate) in a 12-h/12-h light/dark cycle. For the dietary challenge studies, transgenic (TghPNPLA3-I148I and TghPNPLA3-I148M) and nontransgenic wild type (WT) mice were fed either a high-sucrose (Research Diets, D17070603: 73.5 kcal%, NJ, USA) diet which has been demonstrated to strongly induce hepatic steatosis (but not NASH) as widely reported in the literature including a report on the Pnpla3 I148M knock in mice ^50^ for 4 weeks, or a high-fat, high-fructose, high-cholesterol diet (HFFC) (Research Diets D18021203: 20% calories from protein, 40% calories from fat, 40% calories from carbohydrate and 1% cholesterol, NJ, USA) for 20 weeks. The HFFC diet mimics the modern Western diet and has been widely demonstrated inductive to induce NASH ^51-54^.

#### Glucose and insulin tolerance test

After 16 weeks of HFFC diet feeding, glucose tolerance test (GTT) and insulin tolerance test (ITT) for TghPNPLA3-I148I, TghPNPLA3-I148M, and WT were performed by intraperitoneal (i.p.) injection of 2 g/kg glucose ((Millipore-Sigma, MO, USA) or 0.75 unit/kg insulin (Lily, IN, USA). All of the mice were fasted for 5 hours for GTT or 4 hours for ITT prior to intraperitoneal injection. Plasma glucose levels were determined before the injection of glucose or insulin and at 15, 30, 60 and 120 min after the injection by a Contour glucometer (Bayer HealthCare, IN, USA).

#### Glycemic and lipids profile and fat composition measurement

Fasting blood samples were collected biweekly after fasting for 4 h. A 16-hour fasting was performed for blood glucose measurement at the 16th week for the mice fed with the HFFC diet. Plasma glucose levels were measured by a Contour glucometer. The fasting plasma insulin concentration were determined by a mouse ultrasensitive insulin ELISA kit (ALPCO, NH, USA) according to the manufacturer’s protocol.

Hepatic lipid was extracted as previously described ^55^. Total cholesterol and triglycerides concentrations in hepatic or plasma were measured according to manufacturer instruction from commercial assay kits (Wako Chemicals, VA, USA), respectively. Mouse body weights were measured weekly. Total body fat mass and lean mass were evaluated using magnetic resonance imaging (MRI) with an EchoMRI 500-Analyzer (EchoMRI, TX, USA) at the Islet and Physiology Core of Indiana University Center for Diabetes and Metabolic Diseases. The tests were performed at week 20 prior to mice sacrifice. At the end of treatment, mice were sacrificed, and blood, liver, and fat tissue were collected for various analyses.

#### Histological analysis

The liver specimens were fixed in 10% formalin solution and routinely processed for paraffin-embedding, sectioning (5 μm thickness) and hematoxylin and eosin (H&E) staining were performed at the Histology Core of Indiana University School of Medicine following the standard protocol. For Sirius red staining, sections were incubated with a 0.1% direct red 80 (Sigma-Aldrich, MO, USA) plus 0.1% fast green FCF (Fisher Scientific, MA, USA) solution dissolved in saturated aqueous picric acid (1.2% picric acid in water) for 1 hour at room temperature, dehydrated, and mounted with a Permount™ mounting medium (Fisher Scientific, MA, USA). The images of H&E and Sirius red staining were captured using a Leica DM750 microscope equipped with an EC3 digital camera (100× magnifications). The lipid droplets area and Sirius red positive area were quantified using ImageJ software.

#### Immunofluorescence (IF)

Sections were deparaffinized, rehydrated by washes in graded alcohols (ethanol 100%, 95%, 70%, 50%) and distilled water, and retrieved in 0.01 M sodium citrate buffer (10 mM Sodium Citrate, 0.05% Tween 20, pH 6.0) for 20 minutes at 98°C by microwave heating. Sections were blocked with blocking buffer (10% normal serum, 1% bovine serum albumin (BSA) and 0.1% Triton X-100 in PBS) for 2 h at room temperature and incubated overnight at 4°C with rabbit polyclonal anti-myeloperoxidase (MPO, 1:100, Invitrogen, CA, USA) or rabbit monoclonal anti-F4/80 (1:100, Invitrogen, CA, USA) antibody, respectively. After washing with 0.025% Triton X-100 in PBS, sections were incubation with Alexa-Fluor-488 secondary antibodies (Molecular Probes, OR, USA) for 1 h at room temperature, and then mounted with VECTASHIELD® antifade mounting media with DAPI (Vector Laboratories, CA, USA). Immunofluorescence images were captured by a ZEISS fluorescence microscope using an AxionVison Rel 4.8 software (200× magnification). The positive cells were counted in randomly selected fields (five fields per section).

#### Statistical analyses

Two-way repeated measures ANOVA was used to assess the effects of genotype, time, and genotype-time interaction on body weight, glucose, insulin, cholesterol, and triglycerides levels. Tukey’s multiple comparisons test was used to determine the significance of pair-wise comparisons at each time point. Tukey corrected *p* value<0.05 was considered significant. All the statistical analyses for animal experiments were performed using GraphPad Prism Version 6.00 (GraphPad Software, CA, USA)

## RESULTS

### Study overview

The design of this study consists of three steps (**Figure 1**). We first aimed to explore whether NAFLD (both CT measured hepatic steatosis and biopsy-proven histologic NASH) casually affects T2D, obesity and their related intermediate traits. To this end, we used the summary GWAS data for CT scan-measured steatosis (GOLD) as well as the histologic NASH progression (NASH CRN/MIGen) presented in Speliotes et al.^18^, which is thus far the largest GWAS comprehensively covering multiple phenotypes of NAFLD. We also use the summary GWAS meta-analysis data for T2D (DIAGRAM) ^19^, obesity (GIANT) ^26^, glycemic (MAGIC) ^20-25^ and lipid (GLGC) ^27^ traits as outcomes (Step 1), which are the largest-to-date GWAS data on these phenotypes. The causal role of NAFLD on T2D and obesity was conducted by using two well-established NAFLD-associated polymorphisms in PNPLA3 and TM6SF2/NCAN loci as a proxy for hepatic steatosis and NASH progression. We further performed a GWAS on “fatty liver disease” in UK biobank. Using the aforementioned summary level data of DIAGRAM and GIANT, we then investigated the reverse relationships, i.e. whether T2D or obesity causally affect NAFLD risk in the UK Biobank samples (Step 2). Finally, we constructed a transgenic mouse model expressing human PNPLA3-I148I or PNPLA3-I148M isoform to test the causal effects of hepatic steatosis on T2D and obesity (Step 3). To test the effect of steatosis and NASH disease progression on the susceptibility to T2D and obesity, we used a high sucrose diet (HSD) known to induce steatosis in PNPLA3148M mice ^50^, as well as a high fat, high fructose and high cholesterol diet (HFFC) known to induce NASH in mice 51-54. A schematic representation of the three assumptions for an MR analysis was shown in **Figure 2A**, and the MR methods and heterogeneity tests used in the study were listed in **Figure 2B**.

**Figure 2.**
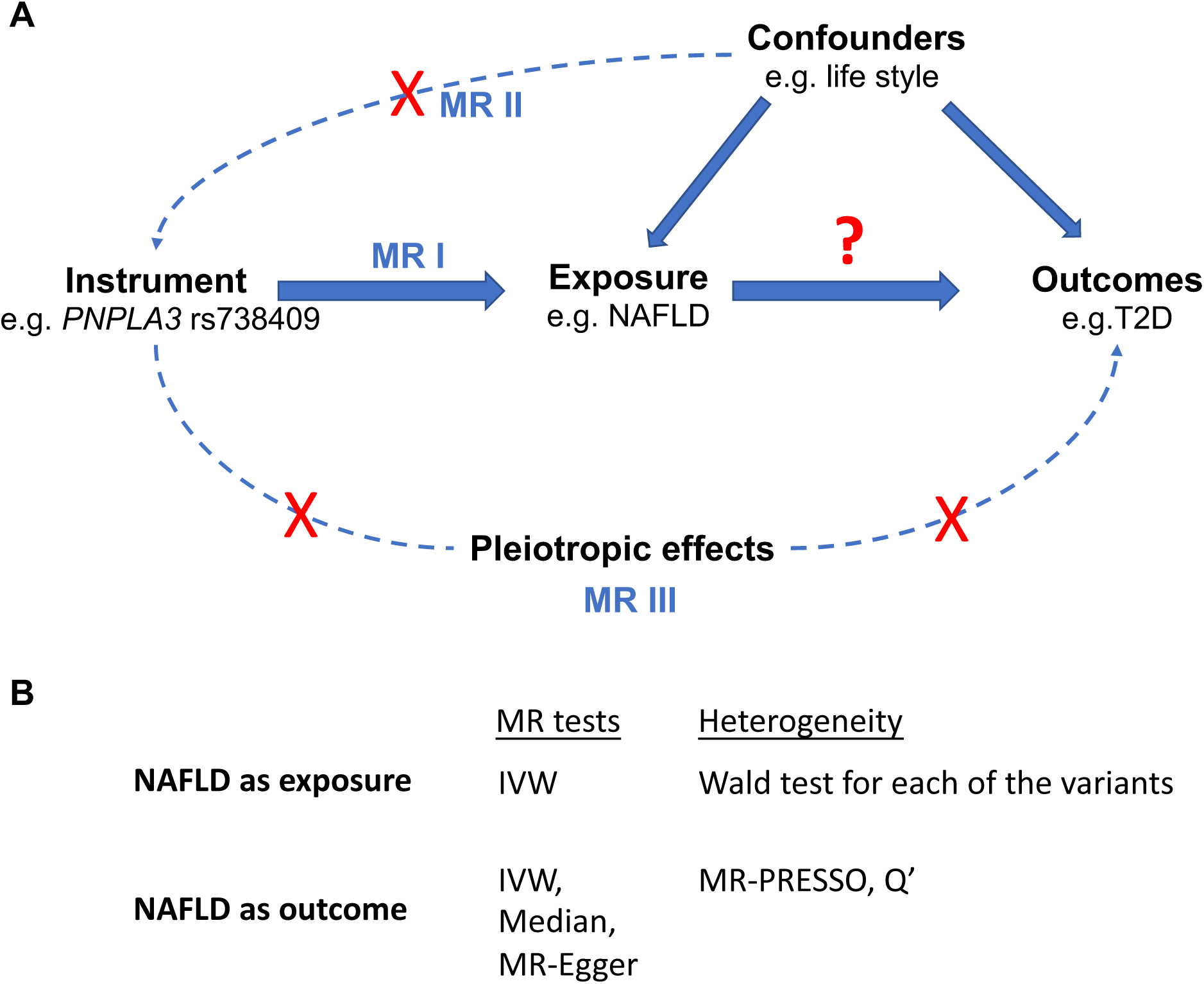
Diagram of the two-sample MR design, three assumptions, and methods used. (A) MR analysis was used to explore the causal relationships among NAFLD, T2D and obesity. Three assumptions of the genetic instrumental variable are as follows: 1) The genetic variant (e.g. PNPLA3 rs738409) is robustly associated with the exposure of interest (e.g. NAFLD), 2) the genetic variant is not associated with confounding factors (e.g. life style), and 3) there is no alternative pathway through which the genetic variant affects the outcomes (e.g. T2D) other than via the exposure. (B) MR methods and heterogeneity tests used in the study. IVW: inverse variance weighted; Median: weighted median estimator; MR-PRESSO: MR pleiotropy residual sum and outlier; Q’: Q’ statistics with modified second order weights;

### The causal effect of NAFLD on T2D risk and glycemic traits

Using two well-established NAFLD-associated variants in PNPLA3 (rs738409) and TM6SF2/NCAN (rs2228603) gene loci as the genetic predictors of steatosis and histologic NASH progression, we tested the causal effect of NAFLD on T2D and glycemic traits in the latest publicly available GWAS data. As listed in **Table 1**, we observed a significant association between genetically instrumented hepatic steatosis and T2D risk, in which a one-standard deviation (SD) increase in CT measured hepatic steatosis caused a 30% increased risk of T2D (OR: 1.3, 95% CI: [1.2, 1.4], *p*=8.3e-14). As for glycemic traits, we detected nominal associations of steatosis with increased fasting glucose (β: 0.026 mmol/L, 95% CI: [8.5e-5, 0.051], *p*=0.049), fasting insulin (β: 0.025 pmol/L, 95% CI: [0.0035, 0.046], *p*=0.022), and insulin resistance (HOMA-IR) (β: 0.03 (mU/L)*(mmol/L), 95% CI: [−0.0016, 0.061], *p*=0.063) levels. However, these results were not significant after adjusted with Bonferroni correction (*p*<2.3e-3, correction for 22 traits). Other tested glycemic traits including HbA1c, fasting proinsulin, 2-h glucose, HOMA-IR, HOMA-B, and seven insulin secretion and action indices during OGTT did not show any significance.

**Table 1.** MR estimate with NAFLD as exposure

| T2D and glycemic traits | Hepatic steatosis (per SD) |  | Histologic NAFLD (per logOR) |  |
| --- | --- | --- | --- | --- |
|  | Effect (95% CI) | p | Effect (95% CI) | p |
| T2D (OR) | 1.30 (1.21, 1.39) | **8.30E-14** | 1.06 (1.03, 1.09) | **2.80E-04** |
| HbA1c (%) | -0.0072 (-0.025, 0.01) | 4.20E-01 | -0.0025 (-0.0074, 0.0024) | 3.10E-01 |
| Fasting glucose (mmol/L) | 0.026 (8.5E-05, 0.051) | 4.90E-02 | 0.0032 (-0.0031, 0.0096) | 3.20E-01 |
| Fasting insulin (pmol/L) | 0.025 (0.0035, 0.046) | 2.20E-02 | 0.0064 (0.00034, 0.012) | 3.80E-02 |
| Fasting proinsulin (pmol/L) | 0.032 (-0.026, 0.089) | 2.80E-01 | 0.0051 (-0.0091, 0.019) | 4.80E-01 |
| 2h glucose (mmol/L) | -0.066 (-0.21, 0.079) | 3.70E-01 | -0.015 (-0.051, 0.021) | 4.10E-01 |
| HOMA-IR ((mU/L)*(mmol/L)) | 0.03 (-0.0016, 0.061) | 6.30E-02 | 0.0066 (-0.0016, 0.015) | 1.10E-01 |
| HOMA-B ((mU/L)/(mmol/L)) | 0.007 (-0.019, 0.033) | 5.90E-01 | 0.0011 (-0.0052, 0.0074) | 7.30E-01 |
| AUCins (mU*min/L) | -0.012 (-0.18, 0.16) | 8.90E-01 | -0.0043 (-0.046, 0.038) | 8.40E-01 |
| AUCins/AUCgluc (mU/mmol) | -0.01 (-0.18, 0.16) | 9.10E-01 | -0.0037 (-0.046, 0.038) | 8.60E-01 |
| Incre30 (mU/L) | -0.0021 (-0.17, 0.16) | 9.80E-01 | -0.00033 (-0.041, 0.04) | 9.90E-01 |
| Ins30adjBMI | 0.017 (-0.15, 0.19) | 8.40E-01 | 0.00083 (-0.041, 0.043) | 9.70E-01 |
| ISI (mg/dL) | 0.011 (-0.18, 0.2) | 9.10E-01 | 0.017 (-0.032, 0.065) | 5.00E-01 |
| CIRadjBMI | -0.0055 (-0.18, 0.17) | 9.50E-01 | -0.0034 (-0.045, 0.039) | 8.80E-01 |
| DI | 0.078 (-0.082, 0.24) | 3.40E-01 | 0.022 (-0.018, 0.063) | 2.80E-01 |
| **Obesity traits** |  |  |  |  |
| BMI (SD) | -0.027 (-0.041, -0.013) | **1.30E-04** | -0.0071 (-0.012, -0.0023) | 3.40E-03 |
| WHR (SD) | 0.021 (0.0062, 0.036) | 5.40E-03 | 0.0029 (-0.00091, 0.0067) | 1.30E-01 |
| WHRadjBMI (SD) | 0.039 (0.023, 0.054) | **8.20E-07** | 0.0072 (0.0022, 0.012) | 4.50E-03 |
| HDL* (SD) | -0.06 (-0.1, -0.016) | 8.20E-03 | -0.013 (-0.025, -0.0014) | 2.80E-02 |
| LDL* (SD) | -0.058 (-0.11, -0.0097) | 1.90E-02 | -0.013 (-0.025, -4E-04) | 4.30E-02 |
| TC* (SD) | -0.084 (-0.13, -0.036) | **6.80E-04** | -0.019 (-0.033, -0.0044) | 9.90E-03 |
| TG* (SD) | 0.02 (-0.023, 0.062) | 3.60E-01 | 0.0043 (-0.0052, 0.014) | 3.80E-01 |

Effect was estimated by a combined genetic vector of PNPLA3 and TM6SF2(NCAN) variants through inverse variance weighted (IVW) method. The F statistics of the genetic predictors of hepatic steatosis and histologic NAFLD are 110 and 10, respectively. Significant results at the Bonferroni-adjusted level of significance ( $p < 0.05/22 = 2.3 \times 10^{-3}$ ) were highlighted in bold.
\*estimate was made using PNPLA3 variant only due to the pleiotropic effect of TM6SF2(NCAN)
T2D: type 2 diabetes; HOMA-IR: The homeostatic model assessment (HOMA) insulin resistance; HOMA-B: HOMA beta cell function; AUCins: area under the curve (AUC) of insulin levels during oral glucose tolerance test (OGTT); AUCins/AUCgluc: ratio of AUC insulin and AUC glucose; Incre30: incremental insulin at 30 min; Ins30adjBMI: Insulin response to glucose during the first 30 min adjusted for BMI; ISI: insulin sensitivity index; CIRadjBMI: Corrected Insulin Response adjusted for ISI; DI: disposition index; BMI: body mass index; WHR: waist-hip ratio; WHRadjBMI: WHR adjusted for BMI; HDL: high-density lipoprotein cholesterol; LDL: low-density lipoprotein cholesterol; TC: total cholesterol; TG: triglycerides; OR: odds ratio; CI: confidence interval; SD: standard deviation.

We then tested if genetically increased risk for the disease progression of NASH also has causal effect on T2D susceptibility and glycemic traits. Consistent with the result of hepatic steatosis, there was evidence of a significant effect of histologically characterized NAFLD severity on an increased risk of T2D (OR: 1.06, 95% CI: [1.03, 1.09], *p*=2.8e-4). No significant causal relationship was found between genetically driven NAFLD and glycemic traits except a nominal association with increased fasting insulin levels (β: 0.0064 pmol/L, 95% CI: [0.00034, 0.012], *p*=0.038) (**Table 1**).

In order to avoid potential biases on the selection of genetic instrument, a post-hoc analysis by separately examining the causal role of PNPLA3 and TM6SF2/NCAN polymorphisms was also performed and generated similar results (**Supplement Table 4**).

### The causal effect of NAFLD on obesity

While obesity is a well-known risk factor for NAFLD, the reverse relationship, i.e. the causal effect of NAFLD on obesity, has not been explored before. Using the same genetic predictors of steatosis and histologic NAFLD, we implemented MR and observed a significant causal association of a one-SD increase in hepatic fat with a 0.027-SD decrease in BMI (β:-0.027, 95%CI: [−0.043, −0.013], *p*=1.3e-4), but a 0.039-SD increase in WHRadjBMI (WHR adjusted for BMI) (β:0.039, 95%CI: [0.023, 0.054], *p*=8.2e-7), an established marker for abdominal or central obesity. Similar relationships were found between genetically raised histologic NAFLD with BMI (β:-0.0071, 95%CI: [−0.012, −0.0023], *p*=3.4e-3) and WHRadjBMI (β:0.0072, 95%CI: [0.0022, 0.012], *p*=4.5e-3). Taken together, our analyses suggested a consistent negative causal relationship between NAFLD and overall obesity (proxied by BMI), but a positive correlation with central or visceral obesity (proxied by WHRadjBMI).

We next investigated the causal effect of NAFLD on blood lipid levels including HDL, LDL, total cholesterol (TC), and triglycerides (TG). Since TM6SF2/NCAN plays a role in plasma lipid regulation and might lead to the invalidity of the MR assumptions, we used PNPLA3 rs738409 variant only with regard to estimating the causal effect on blood lipid profile. Our analyses demonstrated a significant negative correlation between genetically raised hepatic fat and TC levels (β:-0.084 SD, 95%CI: [−0.13, −0.036], *p*=6.8e-4). This negative relationship also exists for histologic NAFLD (β:-0.019 SD, 95%CI: [−0.033, −0.0044], *p*=9.9e-3). No significant causal relationship was found with other lipids (**Table 1**).

### Reverse MR investigating the causal effects of T2D, obesity, and their related secondary traits on NAFLD

To understand the causal relationships among the three diseases, we implemented MR analyses to test the existence of the reverse or bidirectional causal relationships between T2D or obesity and NAFLD. We used the data from our newly performed GWAS on FLD as an outcome, while the DIAGRAM and GIANT summary data for T2D and obesity as predictor, respectively. We found that genetic predictors of T2D exert positive effects on NAFLD (OR: 1.1, 95% CI: [1.0, 1.2], *p*=1.67e-3) without evidence of significant heterogeneity (*P*_MR-PRESSOGlobal_=0.31, P_modified Q_=0.33) after removing outlier variants identified by MR-PRESSO (**Table 2**). Initial MR estimates without removing outliers were shown in **Supplemental Table 5**. Consistent with the previous MR estimate^17^, we found BMI causally increased the NAFLD risk (OR: 2.3 95% CI: [2.0, 2.7], *p*=1.4e-25), but with remaining heterogeneity (*P*_MR-PRESSOGlobal_<2.5e-3, P_modified Q_=3.9e-03) after removing outliers. BMI adjusted WHR also significantly aggravated NAFLD risk (OR: 1.5 95% CI: [1.3, 1.8], *p*=1.1e-06) without evident heterogeneity (*P*_MR-PRESSOGlobal_=0.46, P_modified Q_=0.45).

**Table 2.**
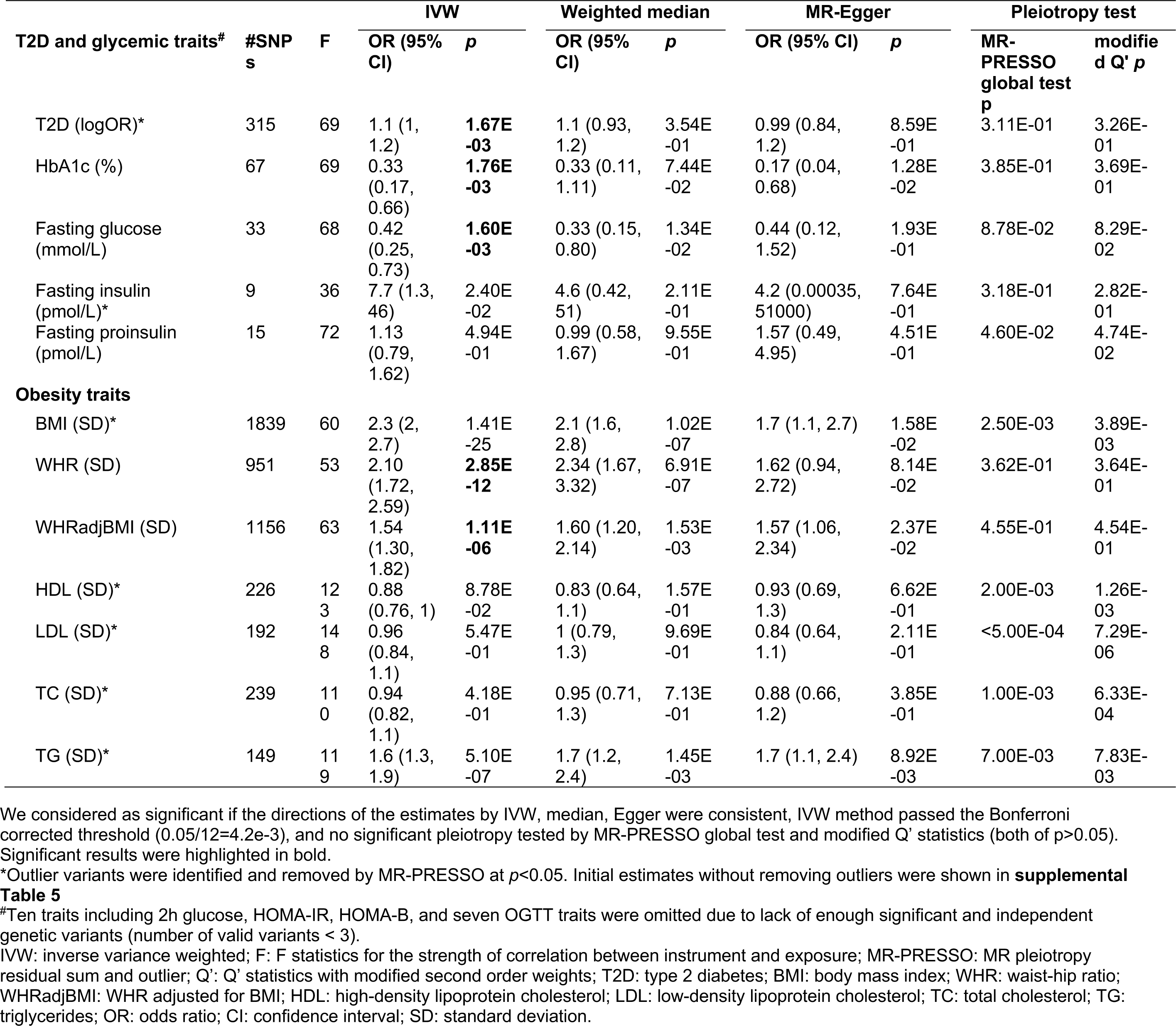
MR estimate with NAFLD as outcome

### Transgenic mice study on the relationship between NAFLD and susceptibility to T2D and obesity

To further examine the causal effect of NAFLD on T2D and obesity, we set out to induce hepatosteatosis and NASH using transgenic animal models expressing human PNPLA3 isoforms, and during which, to observe the development of T2D and obesity phenotypes. To do so, we constructed mice models transduced with the bacterial artificial chromosome (BAC)-containing the PNPLA3-I148I isoform or that was engineered to the PNPLA3-I148M isoform. The mice were then fed with a previously established high sucrose diet (HSD) for 4 weeks to induce hepatosteatosis^50^. To examine the effect of NASH progression on T2D or obesity phenotypes, we also fed the mice with the “Western diet” characterized with high fat, high fructose and high cholesterol (HFFC) for 20 weeks, which has been an established NASH-inductive diet as demonstrated before^51-54^.

After 4 weeks of HSD diet, the TghPNPLA3-I148M mice developed severe hepatosteatosis as compared to their TghPNPLA3-I148I littermates or the non-transgenic controls, characterized with significantly increased lipid droplets formation in the liver (Supplement **Figure S1A**) as well as hepatic triglycerides (TG) accumulation (Supplement **Figure S1B**). Meanwhile, as compared to the TghPNPLA3-I148I controls, the TghPNPLA3-I148M mice also demonstrated a trend of increased circulating glucose, but the insulin level remained unchanged (**Figure 3A, B**). After 4 weeks of HSD diet feeding, the TghPNPLA3-I148M mice demonstrated no significant change in total body weight (**Figure 3C**), but a marginal trend to a reduced total circulating cholesterol (**Figure 3D**, *p*=0.038), but not the circulating TG levels (**Figure 3E**), as compared to their TghPNPLA3-I148I littermates.

**Figure 3.**
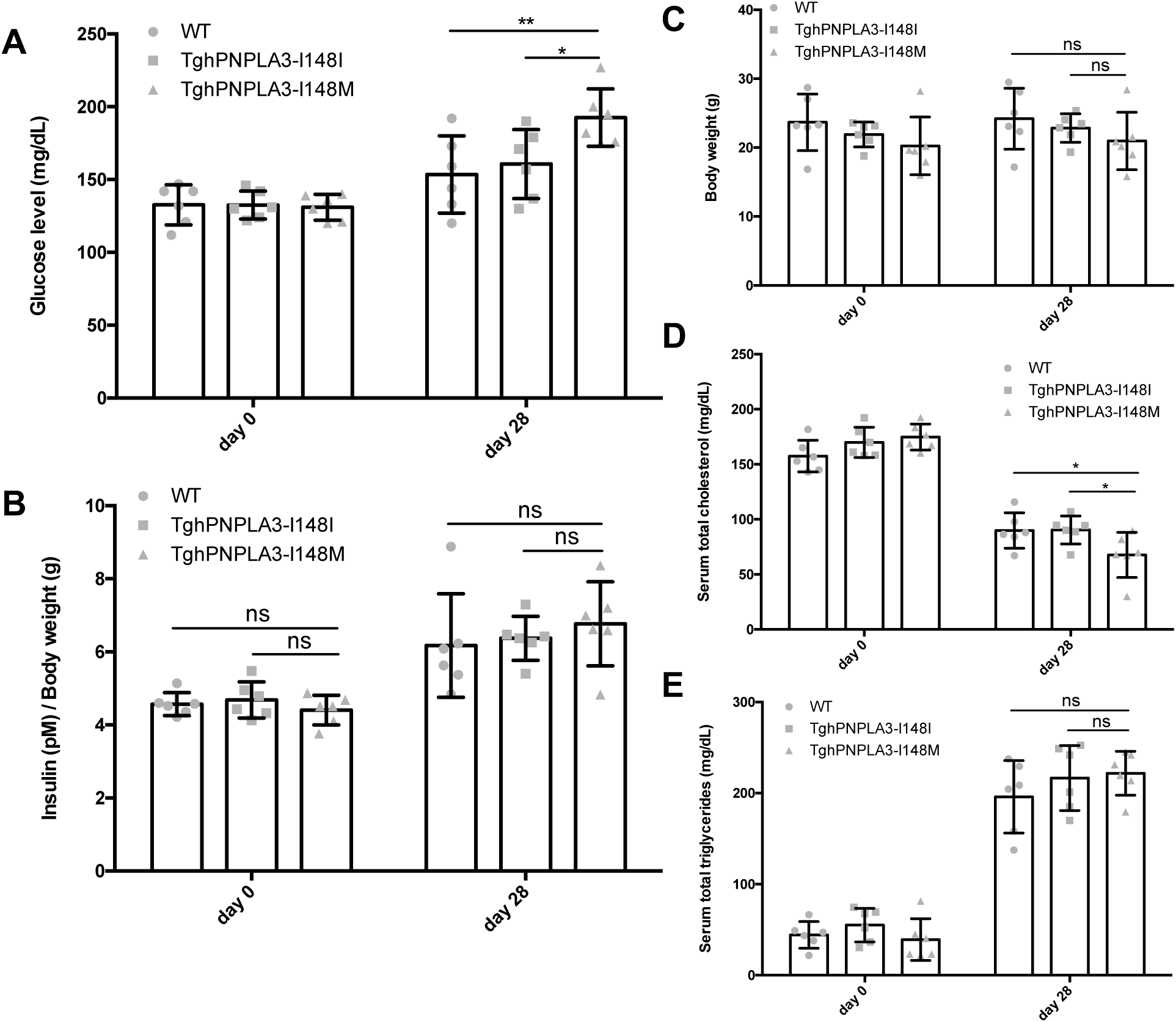
Effect of PNPLA3 I148M mutant on T2D and obesity with an HSD diet. (A) Glucose levels of TghPNPLA3-I148I, TghPNPLA3-I148M, and non-transgenic wide type mice fed with a high sucrose diet (HSD) for 4 weeks. (B) Insulin levels. We normalized the insulin level to the body weight since the age of the TghPNPLA3-I148M group is younger than the TgHPNPLA3-I148I and non-transgenic control groups. We found that the insulin level, especially at the baseline, is highly correlated with body weight (**Supplemental Figure 2A**). The non-normalized data were also presented in **Supplemental Figure 2B**. (C) Body weight (D) Serum total cholesterol levels (E) Serum total triglycerides levels of the three groups fed with an HSD diet for 4 weeks. Error bar represents standard deviation (SD). *: Tukey adjusted *p*<0.05; **: Tukey adjusted *p*<0.01, ns: not significant.

To examine the effect of NASH progression on the susceptibility to T2D and obesity, we fed the mice with the NASH-inducing HFFC diet for 20 weeks. The TghPNPLA3-I148M mice developed significantly more severe NAFLD/NASH phenotypes as compared to the TghPNPLA3-I148I littermates, as characterized by increased inflammation and fibrosis (**Supplement Figures S3-5**), confirming that PNPLA3 I148M possesses a strong genetic predisposition to NAFLD and NASH. H&E staining results indicated that all the three groups have developed hepatosteatosis after 20 weeks of HFFC diet (**Supplement Figures S6**).

We next evaluated the effect of PNPLA3 I148M on glucose homeostasis over a 20-week follow-up. As shown in **Figure 4A**, we observed a significant genotype-time interaction on the fasting glucose level (two-way repeated measure ANOVA, *p*<0.0001), suggesting that the effect of PNPLA3 I148M on glucose levels depended on disease progression. At week 16 and 18, the TghPNPLA3-I148M mice displayed significantly higher fasting glucose levels than their TghPNPLA3-I148I littermates (*ρ*=0.0048 and 0.0082, respectively). There is also a significant interaction between genotype and time on fasting insulin levels between the TghPNPLA3 I148I and TghPNPLA3 I148M groups (*ρ*=0.0009). However, there is no significant difference between the TghPNPLA3-I148M and non-transgenic wildtype mice. Beginning at week 12, the insulin levels between the latter two groups remain unchanged (**Figure 4B**). Results of Glucose tolerance test (GTT) showed that TghPNPLA3-I148M mice experienced a reduced clearance of blood glucose as compared to the TghPNPLA3-I148I controls (*ρ*=0.012) (**Figure 4C**). However, we did not observe a significant difference in the response to insulin challenge in the Insulin tolerance test (ITT) (**Figure 4D**).

**Figure 4.**
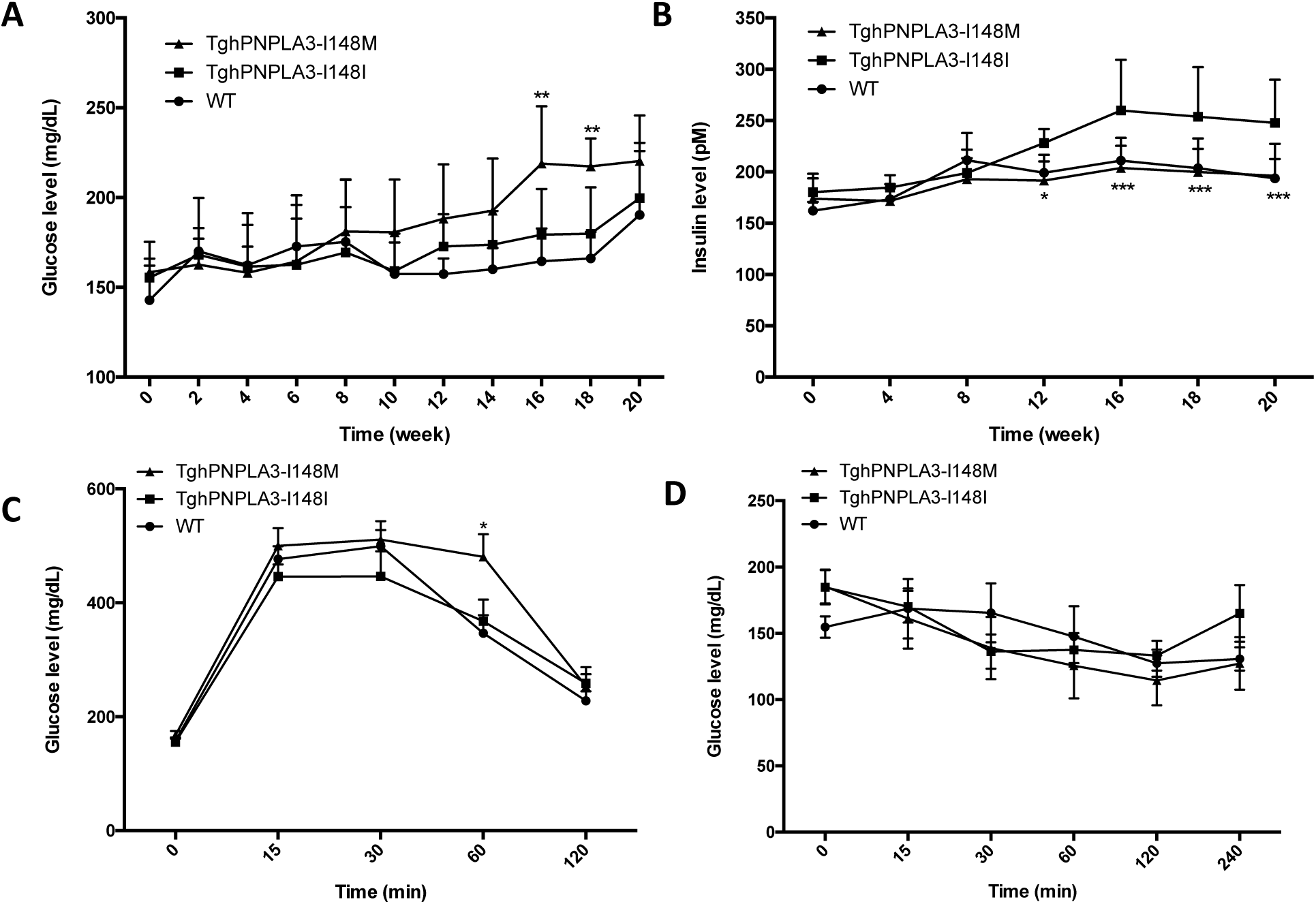
Effect of PNPLA3 I148M mutant on glucose and insulin levels with an HFFC diet. (A) Change in glucose levels over time of TghPNPLA3-I148I, TghPNPLA3-I148M, and non-transgenic wide type mice fed with a high-fat, high-fructose, high-cholesterol (HFFC) diet for 20 weeks. (B) Change in insulin levels for 20 weeks. (C) glucose tolerance test (GTT) and (D) insulin tolerance test (ITT) were performed at the 16th week of HFFC diet feeding. Error bar represents standard deviation (SD). The significance level of the comparison between TghPNPLA3-I148I and TghPNPLA3-I148M was indicated as follows: *: Tukey adjusted *p*<0.05; **: Tukey adjusted *p*<0.01; ***: Tukey adjusted *p*<0.001.

The body weight change of the mice on HFFC diet over time was shown in **Figure 5A**. The TghPNPLA3-I148M mice were significantly lighter than the TghPNPLA3-I148I controls beginning at week 15 (all *ρ*<0.05). Magnetic resonance imaging (MRI) examination at week 20 showed that less total fat was accumulated in the TghPNPLA3-I148M mice than the TghPNPLA3-I148I controls (*ρ*=0.012), while the lean mass of non-fat tissues was not significantly different (**Figure 5B**). Further investigation on the composition of the isolated fat showed that there was a significantly more epididymal white adipose tissue (EWAT) accumulation relative to the total peripheral adipose tissue in the TghPNPLA3-I148M mice than their TghPNPLA3-I148I littermates or non-transgenic controls (*ρ*=0.034 and 0.0015, respectively) (**Figure 5C**). Examining the plasma lipid profile showed a significant decrease in total cholesterol levels beginning at week 16 for TghPNPLA3-I148M mice as compared to the TghPNPLA3-I148I controls (**Figure 5D**). No significant difference in TG levels between the three groups was observed (**Figure 5E**).

**Figure 5.**
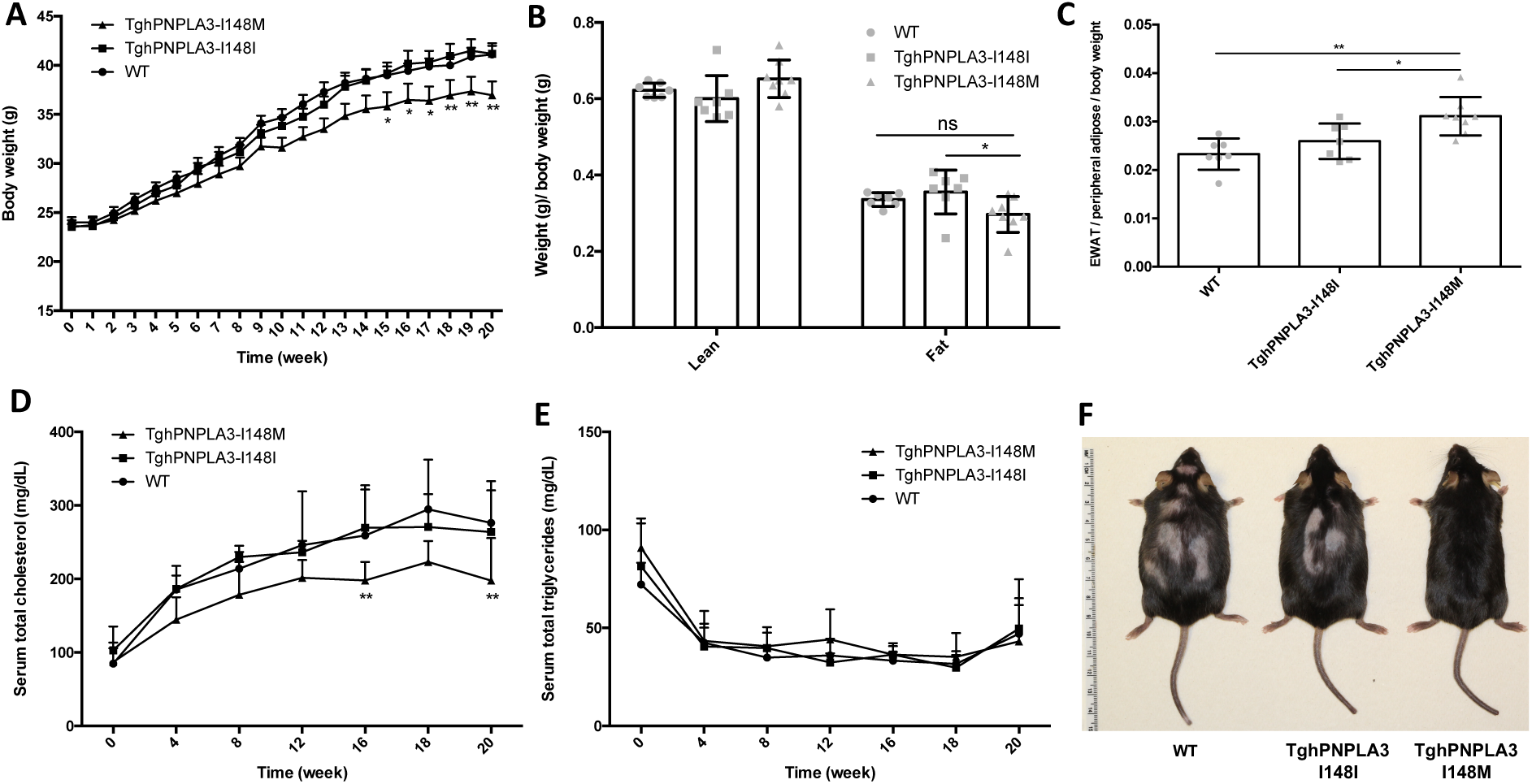
Effect of PNPLA3 I148M mutant on body weight, fat composition, and lipid profiles with an HFFC diet. Change in body weight of TghPNPLA3-I148I, TghPNPLA3-I148M, and non-transgenic wide type mice fed with a high-fat, high-fructose, high-cholesterol (HFFC) diet for 20 weeks. (B) Body composition analysis by magnetic resonance imaging (MRI). Fat tissue weight and non-fat lean mass were normalized by the body weight (C) Epididymal white adipose tissue (EWAT) accumulation at the 20th week of HFFC diet feeding. EWAT accumulation was calculated as weight of EWAT divided the total peripheral adipose tissue weight and then normalized by the body weight. (D) Change in serum total cholesterol levels over 20 weeks (E) Change in serum total triglycerides levels over 20 weeks. (D) The representative image of TghPNPLA3-I148I, TghPNPLA3-I148M, and non-transgenic wide type mouse at the 20th week of HFFC diet feeding. Error bar represents standard deviation (SD). The significance level of the comparison between TghPNPLA3-I148I and TghPNPLA3-I148M was indicated as follows: *: Tukey adjusted P<0.05; **: Tukey adjusted P<0.01; ns: not significant.

## Discussion

This is the first study to delineate the causal inter-relationship between NAFLD, T2D, and obesity using bidirectional MR. This relationship was also experimentally validated using murine models, which further demonstrated a causal relationship between hepatic steatosis or NASH and the susceptibility to T2D and obesity. We found that while genetically driven NAFLD is a causal risk factor for T2D, it protects against overall obesity (indexed by BMI). However, NAFLD SNPs causally increase the risk for central obesity. On the other hand, genetically driven T2D and central or general obesity are all causal for fatty liver disease. Our study thus suggests that genetically driven NAFLD (likely “lean NAFLD”) and the “metabolic NAFLD” attributed to T2D and/or obesity may be different diseases. Similarly, “NAFLD-driven T2D” may be different from T2D caused by other factors as well. Given the active drug development in these areas, it is important to clarify the right disease subtypes to be targeted. Our findings hence have important implications to clinical management, biomedical research and the development of precision medicine for the three diseases.

It has been long regarded that NAFLD and NASH are the central manifestations of T2D and obesity. Due to the lack of data in the natural history of NAFLD/NASH in humans, it remains largely unclear with regard to the causal relationship between NAFLD and T2D or obesity. Without addressing this issue, it is difficult to answer many key questions, e.g. shall we manage the NAFLD in preventing/treating T2D and obesity or vice versa? More importantly, while there is significant heterogeneity in the etiology of each of the three diseases, what would be the right management strategy for the right patient? Delineating this causal relationship is key to precision disease prevention and treatment. With the findings in our study, it is now possible to further dissect the three diseases into important subtypes. First, we observed that genetically driven NAFLD is a causal risk factor for T2D. Our study confirmed the findings of a previous small-scale MR analysis^15^, and is in line with the results from numerous observational studies, including a recent meta-analysis of 19 observational studies with 296,439 individuals which indicated that patients with NAFLD had a two-fold risk of developing T2D than those without NAFLD^3^. More importantly, our animal model study on two different diets also consistently validated that PNPLA3-I148M-driven hepatosteatosis or NASH increases the risk for hyperglycemia. Therefore, based on our study T2D is likely to be divided into at least two subgroups, i.e. a genetically driven NAFLD-associated T2D (“NAFLD driven-T2D”) and T2D due to other etiologies. This suggests that at least a significant proportion of T2D patients or pre-diabetic individuals, i.e. those who carry the PNPLA3-I148M or TM6SF2 variants and develop NAFLD should benefit from an intervention aiming at reducing hepatic steatosis or more advanced liver perturbations. This is particularly consistent with findings of multiple longitudinal studies in Asian populations where nondiabetic individuals with hepatic steatosis at baseline have 2.78-fold increased risk for T2D after 5 years of follow-up; while nondiabetic individuals with steatosis at baseline but not at the follow-up did not demonstrate an increased risk for T2D^56,57^. On the other hand, our findings also demonstrated that T2D is a risk factor for at least a subset of NAFLD, hence, a “T2D-driven NAFLD” that may be a comorbidity of diabetes mellitus. Previous population-based studies suggested that hyperglycemia could be a factor for progression to liver fibrosis, and T2D can enhance the liver stiffness (see review^56^). T2D may also modify the genetic risk of PNPLA3 I148M for NAFLD as well. High glucose level can increase the expression of PNPLA3 via regulating carbohydrate-response element-binding protein (ChREBP)^58^, which is a necessary step for the accumulation of PNPLA3-I148M protein on the surface of lipid droplet in hepatocytes.^50,59,60^ This accumulation further alters the dynamics of hydrolysis of triglycerides and lead to hepatic steatosis^61^. Therefore, individuals who have a high risk for T2D should consider early intervention to further prevent liver injuries.

Our study also revealed an interesting relationship between NAFLD and obesity. Our analysis suggests that genetically driven liver-specific fat storage or deposition may remodel the fat distribution in the whole body. Although epidemiologically obesity is highly correlated with NAFLD, the genetically instrumented NAFLD is actually not a causal risk factor for overall obesity. Rather, it protects against the overall BMI elevation. However, genetically driven NAFLD causally increases risks for central obesity, characterized by an increased waist-to-hip ratio in humans. Our observation is consistent with numerous observational studies on the associations between NAFLD, visceral fat or WHR and BMI^62,63^. Our mouse study also accurately recapitulated these relationships. Interestingly, both MR and animal studies indicated that genetic NAFLD causally leads to decreased total cholesterol (TC) but not triglycerides (TG). Therefore, central obesity, at least in part, may possess a subtype attributed to genetically-driven NAFLD. This is particularly significant, as both TC and TG are generally correlated with visceral fat as demonstrated in many studies^64-66^. The dissociation in our study suggesting that the NAFLD-driven central obesity is likely a unique subtype.

More importantly, our study corroborates the widely discussed hypothesis about “lean NAFLD” and “obese NAFLD”. Both cross-sectional and longitudinal analyses over the past few years have observed significant differences in disease manifestation, progression and clinical outcomes between lean NAFLD and obese NAFLD^67-69^. Our study echoes this observation and indicates that genetically driven NAFLD may be more likely progressed to lean NAFLD (but not necessarily reduces visceral fat accumulation) while does not promote the development of overall obesity. This is also consistent with the observation that PNPLA3 I148M allele is more significantly associated with lean NAFLD^68,69^. Taken together, our study suggests that so-called “lean NAFLD” is not just a phenomenon, but is a disease phenotype based on a certain causal mechanism. On the other hand, our MR analysis demonstrated that both genetically instrumented BMI and WHRadjBMI causally increase the risk for NAFLD, suggesting adiposity can be a risk factor for at least a subset of NAFLD, i.e. “obese NAFLD”, which may synergize the effects of genetic risk factors on the development of NASH or more severe liver injuries^17^, but could also play an independent role. In the latter case, NAFLD is thus more likely a comorbidity of obesity. Therefore, while the management of the lean NAFLD may require more attention to the liver, reducing weight and BMI would be critical to the prevention of obese NAFLD. Collectively, it is now clearer that, depending on the contributing load of genetic risk to each of NAFLD, T2D, and obesity, these diseases should be further dissected into subgroups based on their causal inter-relationship: e.g. NAFLD should be further classified into “genetically-driven NAFLD” or “hepatic NAFLD”, and “metabolic NAFLD” or “systemic NAFLD, while T2D should be also considered at least to be divided into subgroups that are attributed to the hepatic perturbations and extrahepatic modifications (e.g. pancreas), respectively. There should be a NAFLD-driven central obesity as well. The causal relationships among the three diseases and main findings of the study were summarized in **Figure 6**. In future studies, it would be critical to explore how to distinguish these subgroups and identify the high-risk individuals, so that prevention or treatment strategies for each condition can be developed accordingly.

**Figure 6.**
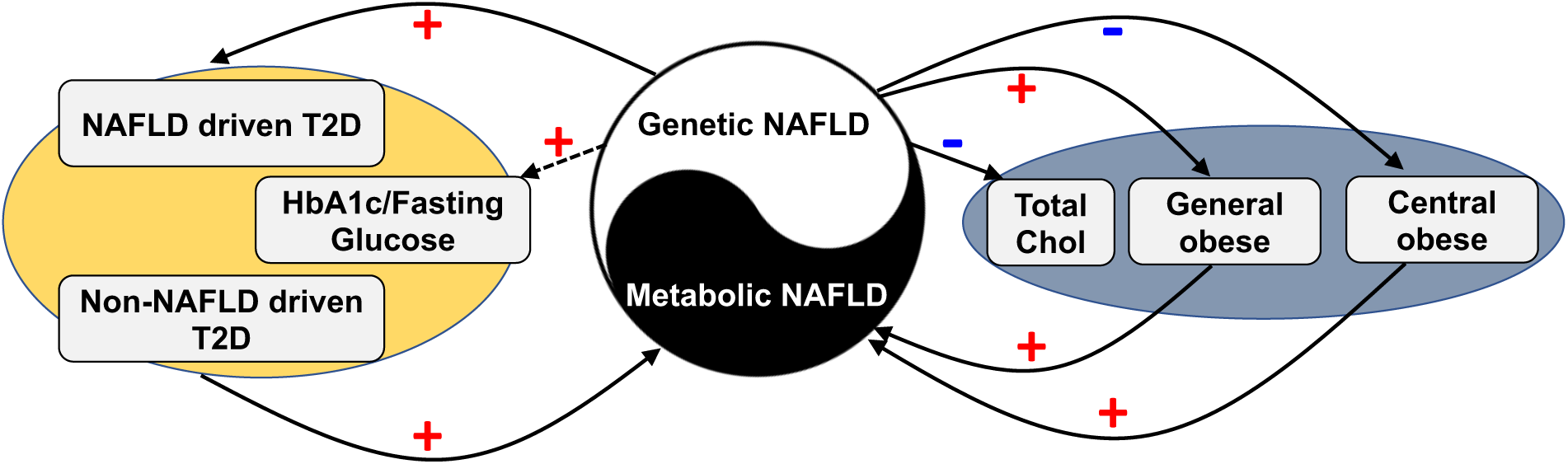
Schematic presentation of the causal relationships among NAFLD, T2D, and obesity. “+”: positive relationship; “-”: negative relationship; dashed line represents the suggestive causal relationship.

Despite the significance of our findings, the detailed molecular mechanism underlying these causal relationships remains to be further investigated. For the causal role of NAFLD in increasing the T2D risk, our MR analysis found a weak causal relationship between genetic NAFLD and fasting glucose and fasting insulin levels. A few hypotheses underlying this NAFLD-T2D relationship were proposed recently, for example, Fetuin-B can cause glucose intolerance, which was deemed as a very convincing mediating mechanism^5,70-73^. In addition, recent MR analyses have demonstrated that central obesity, indexed by increased WHR, is causal to hyperglycemia, whereas the overall obesity is causal to hyperinsulinemia^74,75^, further highlighting the complex interrelationship among these three diseases, as well as their etiological heterogeneity. Whether genetically instrumented NAFLD independently increases the susceptibility to T2D and central obesity or actually affects one after another remains further investigation. Inversely, the mechanisms underlying the causal role of T2D in increasing risk for NAFLD are also incompletely understood, with gut microbiota and expanded, inflamed, dysfunctional adipose tissue as popular hypotheses at present^5^. However, it remains to be further explored whether these potential mechanisms involve the PNPLA3 or TM6SF2 related signaling. Similarly, the causal mechanism underlying the association between genetically driven NAFLD and reduced BMI and total cholesterol but increased abdominal fat accumulation is also largely unclear. Our study warrants continued investigation into these mechanisms.

Our study also has important indications to biomedical research for the three diseases, which is especially pivotal to targets identification and drug development. For instance, the causal pathways leading to “hepatic” or lean NAFLD perhaps are distinct from those of the “metabolic NAFLD”. Selecting the right model would be hence very critical to the success of drug development. Similarly, as mentioned above, determining the correct model for T2D or obesity based on the potential causes may be also key to warrant effective research.

Our study has several limitations. First, we only selected individuals of European descent to avoid the potential confounding effects of population structure. Thus, the findings in our study need to be validated in other ethnic groups; Second, although PNPLA3 and TM6SF2 (NCAN) were two strongest and the most compelling causal variants for NAFLD, the limited number of SNPs prohibits the use of MR-PRESSO and modified Q’ statistics to test the presence of pleiotropic effects. In fact, TM6SF2 variant has been found to be associated with multiple metabolic factors especially lipoprotein lipid profile, therefore may possess potential pleiotropy effects. In addition, TM6SF2 rs58542926 has a relatively lower allele frequency (7% in Europeans) and is also missed from multiple commonly used genotyping platforms. These facts limit its potential as an ideal proxy for an MR analysis. Although our GWAS using the UK Biobank produced more loci that are potentially associated with NAFLD, after a stringent selection, the PNPLA3 and TM6SF2/NCAN loci polymorphisms remain to be the most reliable markers as an instrument for NAFLD. Therefore, it is possible that the causal relationships we have identified in this analysis are only limited to these two genes. In addition, the UK biobank samples only have limited clinical information. We can only focus on the ICD codes for “fatty liver disease” as a phenotype. Although we tried our best to remove other potential confounding factors, e.g. individuals with other known liver disease or hepatitis, the phenotype may not exactly reflect the strictly defined “non-alcoholic fatty liver disease”. However, both PNPLA3 and TM6SF2 loci were successfully identified as top hits in this study (Supplement **Table S6**), suggesting it does reflect the genetic patterns underlying NAFLD. Therefore, the reverse MR to test the causal role of T2D or obesity in increasing risks for NAFLD are likely reliable. Future studies should consider using a large sample set with well-defined NAFLD/NASH phenotypes to further validate these relationships. Moreover, due to the complexity of the biological systems, the existence of the reciprocal feedback loops might mask the truly bidirectional causal relationship. Future studies should consider using alternative approaches such as structural equation modeling to estimate the effect of feedback loops ^76^. Further studies on the contributing load of genetic risk to each of the three diseases and their interactions with the environmental factors are necessary to apply the findings at the individual level.

In summary, our study combined both a bidirectional MR analysis with the largest-to-date sample sets and animal models to delineate the causal relationships between NAFLD, T2D, and obesity. Our findings provided strong evidence for disease subphenotyping and corroborated key hypotheses previously generated from observational studies on these three important diseases. These identifications warranted new directions to the development of precision preventive and therapeutic strategies for the three diseases.

## ACKNOWLEDGEMENTS

This study is supported in part by the NIH/NIDDK grant (R01DK106540) (W.L.), and the start-up fund of the Office of Vice President for Research of Wayne State University (W.L.), NIH R21AA024550 (X.C.D.), R01DK091592 (X.C.D), R56DK091592 (X.C.D), and Indiana Diabetes Research Center grant NIH P30DK097512. We thank all the genetics consortiums for making the GWAS summary data publicly available. We would also like to thank Drs. Anjaneyulu Kowluru, James Granneman, Menghao Huang, as well as Jean-Christophe Rochet and his group for their insightful discussion and comments on this study.

