## Supplemental Figures and Tables for "Mendelian Randomization Analysis Dissects the Relationship between NAFLD, T2D, and Obesity and Provides Implications to Precision Medicine"

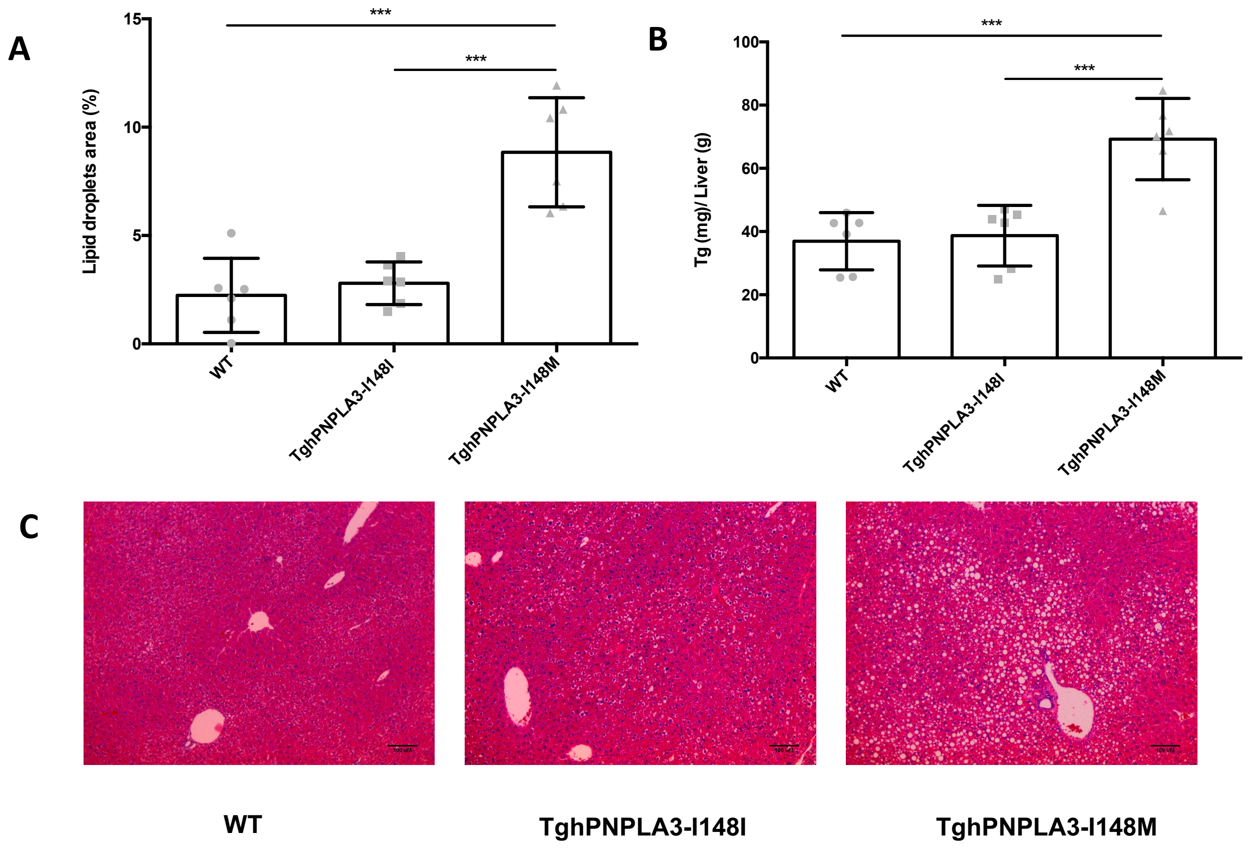


**Supplemental Figure S1. Lipid droplets and TG accumulation of mice fed with an HSD diet for 4 weeks**

(A) Lipid droplets area of TghPNPLA3-I148I, TghPNPLA3-I148M, and non-transgenic wide type mice fed with an HSD diet for 4 weeks. (B) Liver triglycerides levels of the three groups after 4 weeks. (C) H&E staining of liver sections. Error bar represents standard deviation (SD). The significance level of the comparison between TghPNPLA3-I148I and TghPNPLA3-I148M was indicated as follows: *: Tukey adjusted $p<0.05$; **: Tukey adjusted $p<0.01$; ***: Tukey adjusted $p<0.001$.

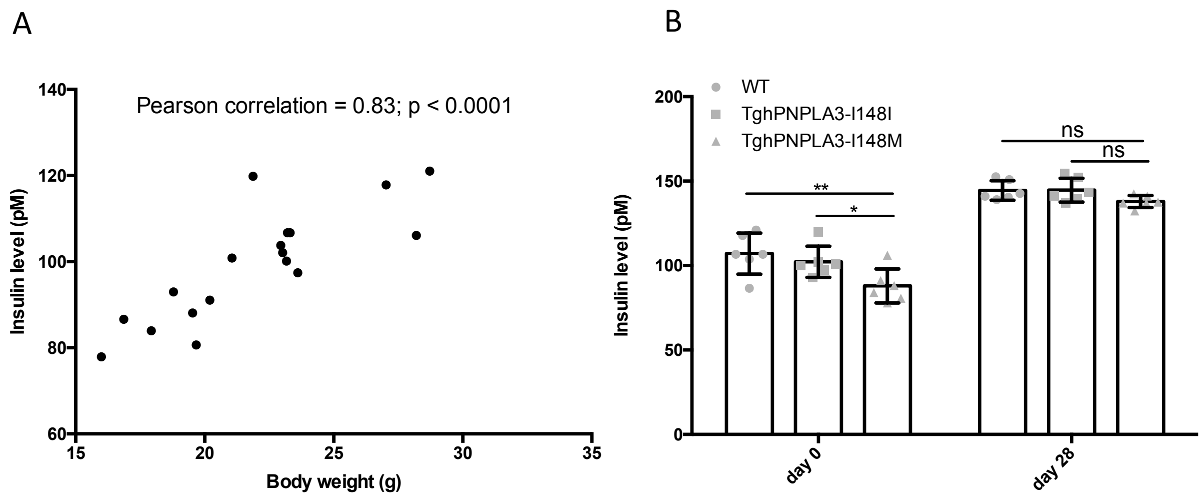


**Supplemental Figure S2. Insulin levels and body weight of the mice fed with an HSD diet**

(A) Pearson correlation between insulin levels and body weight at the baseline of the HSD diet. (B) Insulin levels of TghPNPLA3-I148I, TghPNPLA3-I148M, and non-transgenic wide type mice without normalizing to the body weight. Error bar represents standard deviation (SD). *: Tukey adjusted $p$<0.05; **: Tukey adjusted $p$<0.01; ns: not significant.

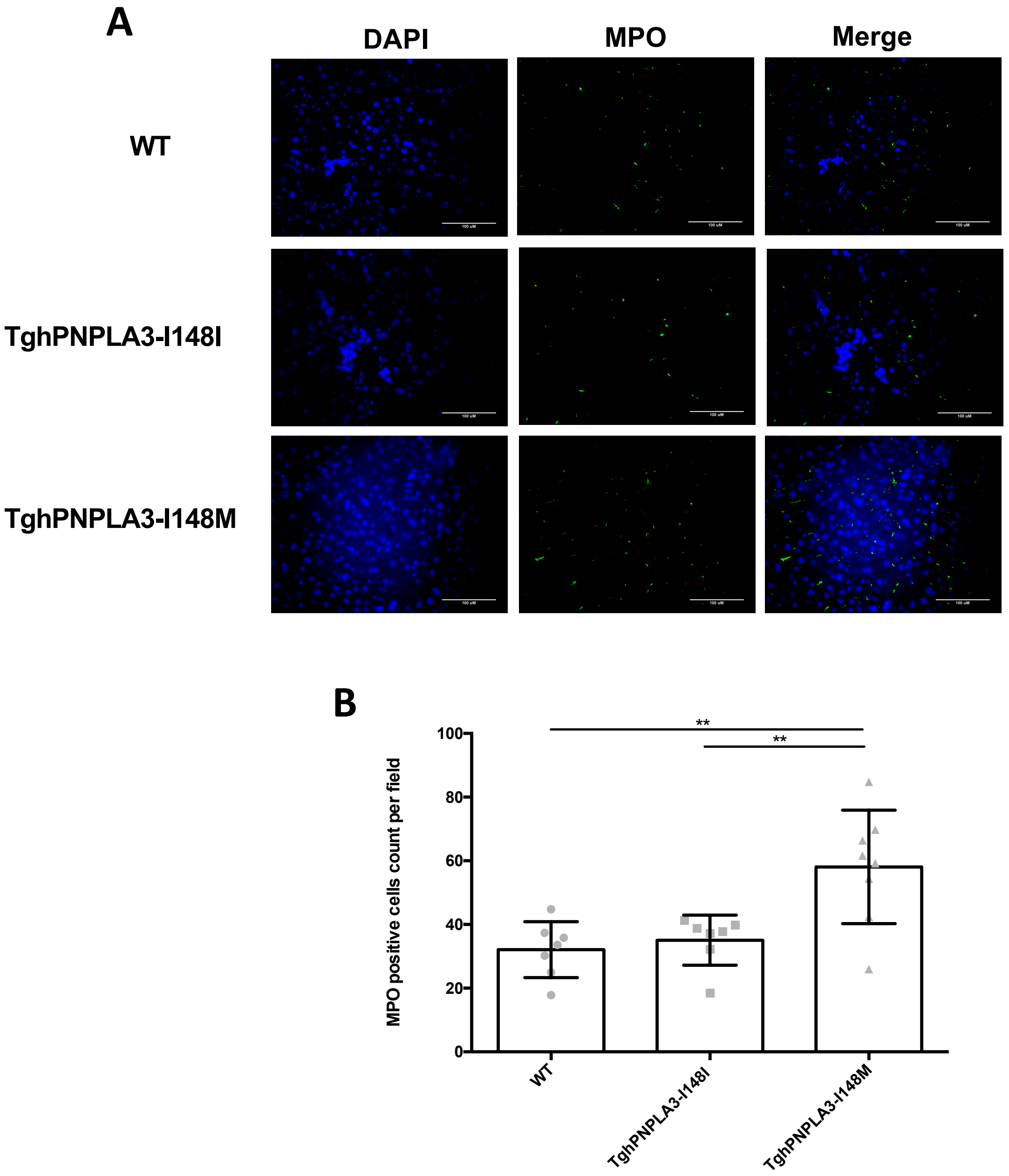


**Supplemental Figure S3. MPO staining of mice fed with an HFFC diet for 20 weeks**

(A) Representative images of MPO immunofluorescence staining. (B) Counts of MPO positive cells in TghPNPLA3-I148I, TghPNPLA3-I148M, and non-transgenic wide type controls. The positive cells were counted in randomly selected fields (five fields per section). Error bar represents standard deviation (SD). *: Tukey adjusted $p$<0.05; ***: Tukey adjusted $p$<0.001.

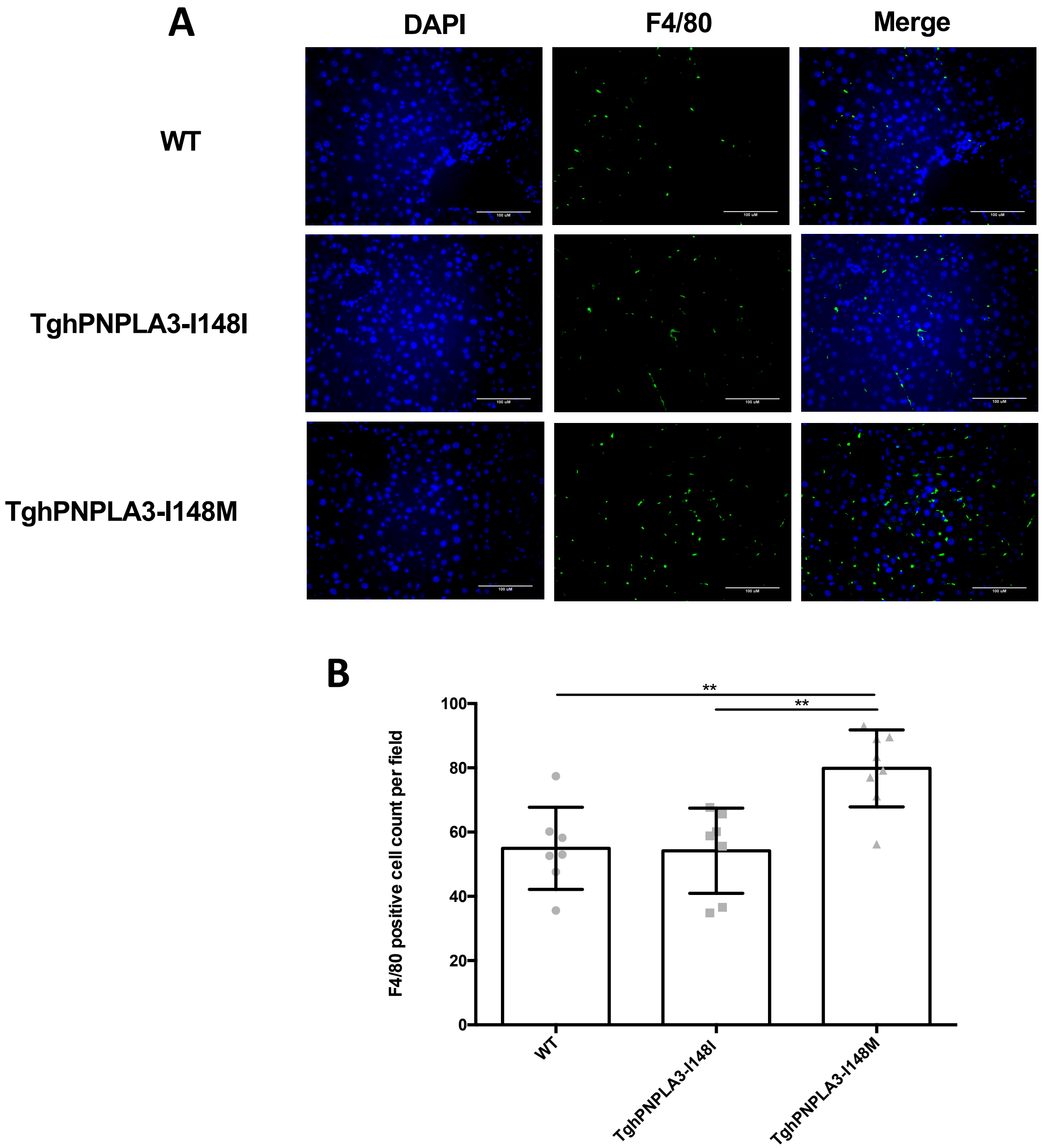


**Supplemental Figure S4. F4/80 staining of mice fed with an HFFC diet for 20 weeks**

(A) Representative images of F4/80 immunofluorescence staining. (B) Counts of F4/80 positive cells in TghPNPLA3-I148I, TghPNPLA3-I148M, and non-transgenic wide type controls. The positive cells were counted in randomly selected fields (five fields per section). Error bar represents standard deviation (SD). *: Tukey adjusted $p<0.05$; ***: Tukey adjusted $p<0.001$.

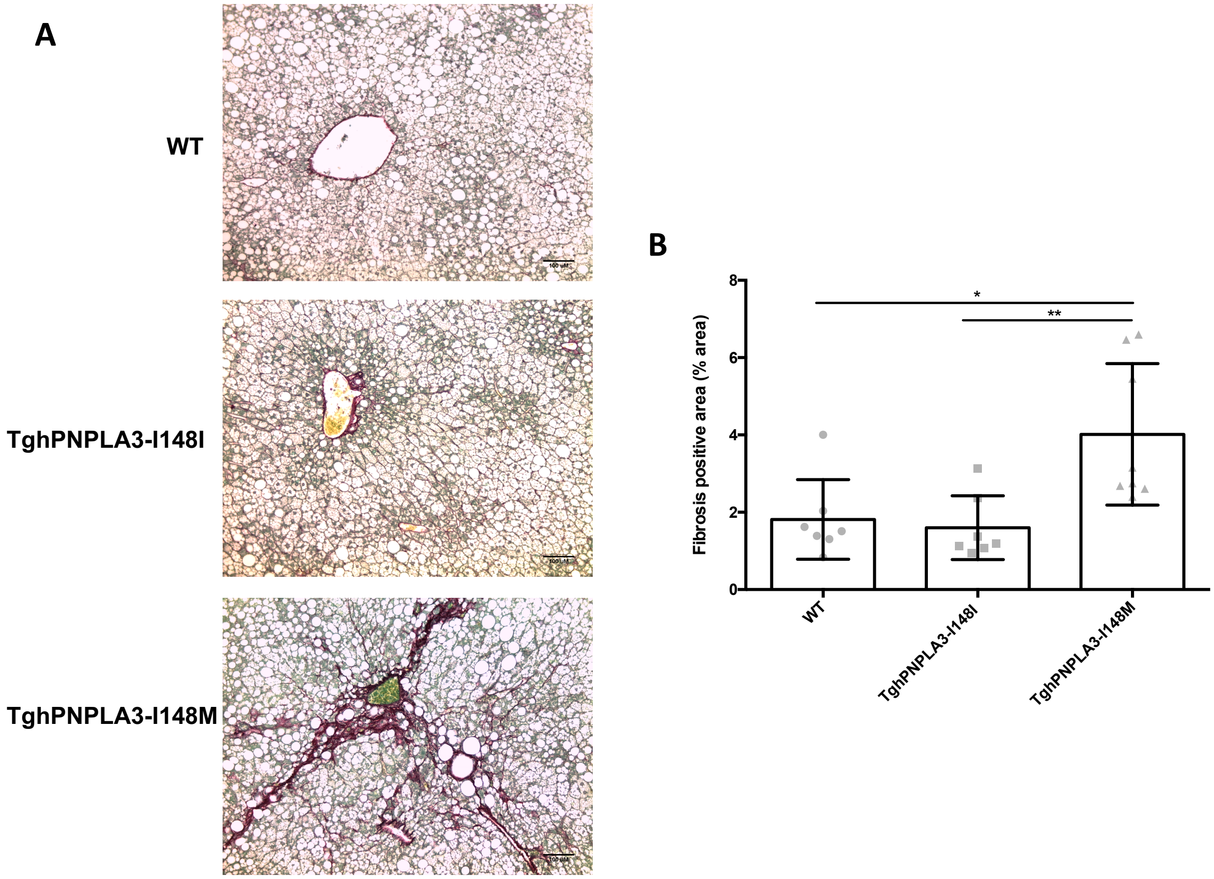


**Supplemental Figure S5. Sirius red staining of mice fed with an HFFC diet for 20 weeks**

(A) Representative images of Sirius red staining. (B) Percentage of fibrosis positive area in TghPNPLA3-I148I, TghPNPLA3-I148M, and non-transgenic wide type mice. Error bar represents standard deviation (SD). *: Tukey adjusted $p<0.05$; **: Tukey adjusted $p<0.01$.

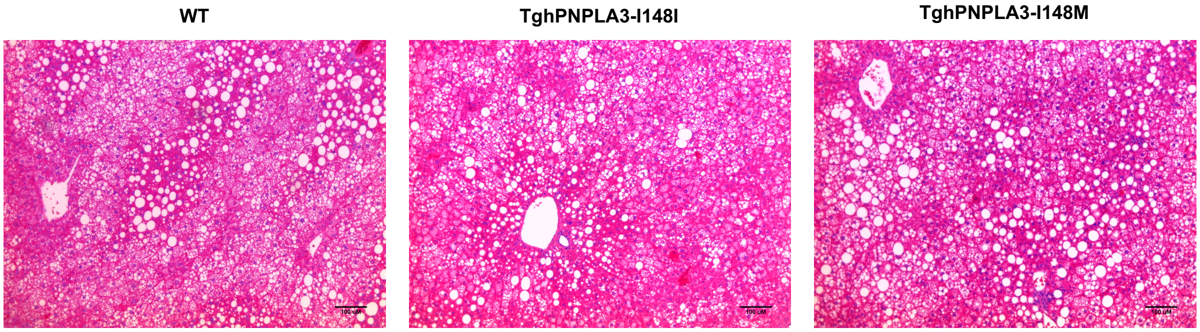


**Supplemental Figure S6. H&E staining of mice fed with an HFFC diet for 20 weeks**

H&E staining of liver sections of TghPNPLA3-I148I, TghPNPLA3-I148M, and non-transgenic wide type mice.

### SUPPLEMENTAL TABLES

**Supplemental Table 1**. Characteristics of the GWAS summary data

| Phenotype | Participants | Data source | Phenotype transformation | Units of effect size | Phenotype description | PubMed ID |
|---|---|---|---|---|---|---|
| **NAFLD** | | | | | | |
| Steatosis | 7,176 | GOLD | inverse normally transformation | SD | Computerized tomography measured hepatic steatosis | 21423719 |
| Histologic NAFLD | 592 cases and 1,405 controls | NASH CRN/MIGen | none | OR | Biopsy-proven NAFLD | 21423719 |
| NAFLD* | 1,122 cases 399,900 controls | UKBB | none | OR | NAFLD from UK Biobank database | Not published |
| **T2D and glycemic traits** | | | | | | |
| T2D | 74,124 cases and 824,006 controls | DIAGRAM | none | OR | Type 2 diabetes adjusted for body mass index (BMI) | 30297969 |
| HbA1c | 123,665 | MAGIC | none | % | Glycated hemoglobin | 28898252 |
| Fasting glucose | 58,074 | MAGIC | none | mmol/L | Fasting glucose adjusted for BMI | 22581228 |
| Fasting insulin | 51,750 | MAGIC | log transformation | pmol/L | Fasting insulin adjusted for BMI | 22581228 |
| Fasting proinsulin | 10,701 | MAGIC | log transformation | pmol/L | Fasting proinsulin adjusted for fasting insulin | 21873549 |
| 2h glucose | 15,234 | MAGIC | none | mmol/L | 2 h glucose levels after an oral glucose challenge adjusted for BMI | 20081857 |
| HOMA-IR | 37,037 | MAGIC | log transformation | original | The homeostatic model assessment (HOMA) insulin resistance | 20081858 |
| HOMA-B | 36,466 | MAGIC | log transformation | original | HOMA beta cell function | 20081858 |
| AUCins | 4,324 | MAGIC | log transformation | mU*min/L | Area under the curve (AUC) of insulin levels during OGTT | 24699409 |

| | | | | | | |
|---|---|---|---|---|---|---|
| AUCins/AUCgluc | 4,213 | MAGIC | log transformation | mU/mmol | Ratio of AUC insulin and AUC glucose | 24699409 |
| Incre30 | 4,447 | MAGIC | log transformation | mU/L | Incremental insulin at 30 min | 24699409 |
| Ins30adjBMI | 4,409 | MAGIC | log transformation | original | Insulin response to glucose during the first 30 min adjusted for BMI =insulin at 30 min/ (glucose at 30 min×BMI) | 24699409 |
| ISI | 4,769 | MAGIC | log transformation | mg/dL | Insulin sensitivity index =10,000/square root (fasting plasma glucose (mg/dl) ×fasting insulin×mean glucose during OGTT (mg/dl) ×mean insulin during OGTT) | 24699409 |
| CIRadjISI | 4,789 | MAGIC | log transformation | original | Corrected Insulin Response =(100 × insulin at 30 min)/ (glucose at 30 min × (glucose at 30 min–3.89)), adjusted for ISI | 24699409 |
| DI | 5,130 | MAGIC | log transformation | original | Disposition index =CIR×ISI | 24699409 |
| **Obesity traits** | | | | | | |
| BMI | 806,834 | GIANT | inverse normally transformation | SD | Body mass index | 30239722 |
| WHR | 697,734 | GIANT | inverse normally transformation | SD | Waist-hip ratio | 30239722 |
| WHRadjBMI | 694,649 | GIANT | inverse normally transformation | SD | Waist-hip ratio adjusted for BMI | 30239722 |
| HDL | 188,577 | GLGC | quantile normalization | SD | Plasma high-density lipoprotein cholesterol | 24097068 |
| LDL | 188,577 | GLGC | quantile normalization | SD | Plasma low-density lipoprotein cholesterol | 24097068 |
| TC | 188,577 | GLGC | quantile normalization | SD | Plasma total cholesterol | 24097068 |
| TG | 188,577 | GLGC | quantile normalization | SD | Plasma triglycerides | 24097068 |

*NAFLD was defined based on ICD-9 571.8 “Other chronic nonalcoholic liver disease”) and ICD- 10 K76.0 [“Fatty (change of) liver, not elsewhere classified”] from inpatient hospital diagnosis within the UK Biobank dataset. Individuals with Hepatitis B or C or with other known liver diseases (e.g. liver transplant, hepatomegaly, jaundice, or abnormal liver function study results) were excluded from the analysis.
GOLD: Genetics of Obesity-related Liver Disease; NASH CRN: NASH Clinical Research Network; MIGen: Myocardial Infarction Genetics consortium; UKBB: UK Biobank; DIAGRAM: DIAbetes Genetics Replication And Meta-analysis; MAGIC: Meta-Analyses of Glucose and Insulin-

related traits Consortium; GIANT: The Genetic Investigation of ANthropometric Traits consortium; GLGC: The Global Lipids Genetics Consortium; OR: odds ratio; SD: standard deviation.

**Supplemental Table 2**. Characteristics of the associations of PNPLA3 rs738409, NCAN rs2228603 with phenotype

| Phenotype | Participants | *PNPLA3* rs738409 G allele | | | *NCAN* rs2228603 T allele | | |
|---|---|---|---|---|---|---|---|
| | | Effect | SE | *p* | Effect | SE | *p* |
| **NAFLD** | | | | | | | |
| Steatosis (SD) | 7,176 | 0.26 | 0.021 | 4.3E-34 | 0.24 | 0.035 | 1.22E-11 |
| Histologic NAFLD (logOR) | 592 cases and 1,405 controls | 1.18 | 0.31 | 3.6E-43 | 0.5 | 0.23 | 5.29E-05 |
| **T2D and glycemic traits** | | | | | | | |
| T2D (OR) | 74,124 cases and 824,006 controls | 0.062 | 0.0089 | 3.5E-12 | 0.086 | 0.014 | 1.30E-09 |
| HbA1c (%) | 123,665 | -0.0037 | 0.003 | 2.2E-01 | 0.0007 | 0.0033 | 8.33E-01 |
| Fasting glucose (mmol/L) | 58,074 | 0.0029 | 0.0038 | 4.4E-01 | 0.02 | 0.0063 | 1.77E-03 |
| Fasting insulin (pmol/L) | 51,750 | 0.0077 | 0.0032 | 1.6E-02 | 0.0024 | 0.0053 | 6.49E-01 |
| Fasting proinsulin (pmol/L) | 10,701 | 0.004 | 0.0087 | 6.4E-01 | 0.02 | 0.014 | 1.44E-01 |
| 2h glucose (mmol/L) | 15,234 | -0.016 | 0.022 | 4.7E-01 | -0.019 | 0.036 | 6.09E-01 |
| HOMA-IR ((mU/L)*(mmol/L)) | 37,037 | 0.007 | 0.0047 | 1.4E-01 | 0.0097 | 0.0081 | 2.33E-01 |
| HOMA-B ((mU/L)/(mmol/L)) | 36,466 | 0.0007 | 0.0039 | 8.5E-01 | 0.0053 | 0.0066 | 4.26E-01 |
| AUCins (mU*min/L) | 4,324 | -0.0066 | 0.026 | 8.00E-01 | 0.0079 | 0.044 | 8.56E-01 |
| AUCins/AUCgluc (mU/mmol) | 4,213 | -0.0057 | 0.026 | 8.30E-01 | 0.0073 | 0.044 | 8.69E-01 |
| Incre30 (mU/L) | 4,447 | -0.00015 | 0.025 | 9.95E-01 | -0.0018 | 0.043 | 9.66E-01 |
| Ins30adjBMI | 4,409 | -0.0027 | 0.026 | 9.16E-01 | 0.026 | 0.043 | 5.50E-01 |
| ISI (mg/dL) | 4,769 | 0.03 | 0.029 | 2.94E-01 | -0.066 | 0.043 | 1.27E-01 |
| CIRadjISI | 4,789 | -0.0062 | 0.026 | 8.09E-01 | 0.013 | 0.043 | 7.60E-01 |
| DI | 5,130 | 0.03 | 0.024 | 2.15E-01 | -0.012 | 0.041 | 7.62E-01 |
| **Obesity traits** | | | | | | | |
| BMI (SD) | 806,834 | -0.0082 | 0.0021 | 7.2E-05 | -0.004 | 0.0031 | 1.98E-01 |
| WHR (SD) | 697,734 | 0.0029 | 0.0022 | 1.8E-01 | 0.0134 | 0.0033 | 3.60E-05 |
| WHRadjBMI (SD) | 694,649 | 0.0078 | 0.0022 | 3.5E-04 | 0.0178 | 0.0033 | 5.95E-08 |
| HDL (SD) | 188,577 | -0.016 | 0.0058 | 9.8E-03 | 0.0054 | 0.0067 | 4.51E-01 |
| LDL (SD) | 188,577 | -0.0151 | 0.0063 | 2.0E-02 | -0.104 | 0.0072 | 4.43E-44 |
| TC (SD) | 188,577 | -0.0219 | 0.0062 | 6.0E-04 | -0.12 | 0.0069 | 1.05E-62 |
| TG (SD) | 188,577 | 0.0051 | 0.0056 | 2.5E-01 | -0.11 | 0.0065 | 1.74E-57 |

T2D: type 2 diabetes; HOMA-IR: The homeostatic model assessment (HOMA) insulin resistance; HOMA-B: HOMA beta cell function; AUCins: area under the curve (AUC) of insulin levels during oral glucose tolerance test (OGTT); AUCins/AUCgluc: ratio of AUC insulin and AUC glucose; Incre30: incremental insulin at 30 min;

Ins30adjBMI: Insulin response to glucose during the first 30 min adjusted for BMI; ISI: insulin sensitivity index; CIRadjBMI: Corrected Insulin Response adjusted for ISI; DI: disposition index; BMI: body mass index; WHR: waist-hip ratio; WHRadjBMI: WHR adjusted for BMI; HDL: high-density lipoprotein cholesterol; LDL: low-density lipoprotein cholesterol; TC: total cholesterol; TG: triglycerides; OR: odds ratio; SD: standard deviation; SE: standard error.

**Supplemental Table 3. Sample overlap**

| Phenotype | Steatosis (GOLD) | | Histologic NAFLD (NASH CRN/MIGen) | | NAFLD (UKBB) | |
|---|---|---|---|---|---|---|
| | **Overlapping cohort** | **Overlapping percentage** | **Overlapping cohort** | **Overlapping percentage** | **Overlapping cohort** | **Overlappin percentage** |
| T2D (DIAGRAM) | FHS | 0.13% | None | None | UKBB | 0.12% |
| HbA1c (MAGIC) | FHS | 1.61% | None | None | None | None |
| Fasting glucose (MAGIC) | FHS | 5.04% | None | None | None | None |
| Fasting insulin (MAGIC) | FHS | 5.67% | None | None | None | None |
| Fasting proinsulin (MAGIC) | FHS | 27.38% | None | None | None | None |
| 2h glucose (MAGIC) | FHS | 17.87% | None | None | None | None |
| HOMA-IR (MAGIC) | FHS | 7.92% | None | None | None | None |
| HOMA-B (MAGIC) | FHS | 8.04% | None | None | None | None |
| Seven oGTT traits (MAGIC) | None | None | None | None | None | None |
| BMI (GIANT) | None | None | None | None | UKBB | 49.70% |
| WHR (GIANT) | None | None | None | None | UKBB | 57.47% |
| WHRadjBMI (GIANT) | None | None | None | None | UKBB | 57.73% |
| Four plasma lipids (GLGC) | Amish | 0.57% | None | None | None | None |

The sample overlapping rate was calculated as the percentage of overlapped samples in the larger dataset. For example, if sample size of study 1 is n1, sample size of study 2 is n2, sample size of the overlapped cohort in study 1 is m1, sample size of the overlapped cohort in study 2 is m2, then the maximum overlapping rate = min(m1,m2) / max(n1, n2).

GOLD: Genetics of Obesity-related Liver Disease; NASH CRN: NASH Clinical Research Network; MIGen: Myocardial Infarction Genetics consortium; UKBB: UK Biobank; DIAGRAM: DIAbetes Genetics Replication And Meta-analysis; MAGIC: Meta-Analyses of Glucose and Insulin-related traits Consortium; GIANT: The Genetic Investigation of ANthropometric Traits consortium; GLGC: The Global Lipids Genetics Consortium; FHS: Framingham Heart Study

**Supplemental Table 4**. Full results of MR estimate with NAFLD as exposure

| Phenotype | Hepatic steatosis (per SD) | | | | | | Histologic NAFLD (per logOR) | | | | | |
|---|---|---|---|---|---|---|---|---|---|---|---|---|
| | *PNPLA3* rs738409 | | *NCAN* rs2228603 | | PNPLA3 rs738409 + NCAN rs2228603 | | *PNPLA3* rs738409 | | *NCAN* rs2228603 | | PNPLA3 rs738409 + NCAN rs2228603 | |
| **T2D and glycemic traits** | **Effect (95% CI)** | ***p*** | **Effect (95% CI)** | ***p*** | **Effect (95% CI)** | ***p*** | **Effect (95% CI)** | ***p*** | **Effect (95% CI)** | ***p*** | **Effect (95% CI)** | ***p*** |
| T2D (OR) | 1.27 (1.17, 1.36) | 1.20E-09 | 1.43 (1.23, 1.68) | 5.20E-06 | 1.30 (1.21, 1.39) | 8.30E-14 | 1.05 (1.02, 1.09) | 9.10E-04 | 1.19 (1.02, 1.39) | 4.30E-02 | 1.06 (1.03, 1.09) | 2.80E-04 |
| HbA1c (%) | -0.014 (-0.037, 0.0085) | 2.20E-01 | 0.0029 (-0.024, 0.03) | 8.30E-01 | -0.0072 (-0.025, 0.01) | 4.20E-01 | -0.0031 (-0.0084, 0.0021) | 2.40E-01 | 0.0014 (-0.026, 0.029) | 8.30E-01 | -0.0025 (-0.0074, 0.0024) | 3.10E-01 |
| Fasting glucose (mmol/L) | 0.011 (-0.017, 0.04) | 4.50E-01 | 0.084 (0.027, 0.14) | 4.00E-03 | 0.026 (8.5e-05, 0.051) | 4.90E-02 | 0.0025 (-0.004, 0.0089) | 4.50E-01 | 0.04 (-0.017, 0.097) | 7.50E-02 | 0.0032 (-0.0031, 0.0096) | 3.20E-01 |
| Fasting insulin (pmol/L) | 0.03 (0.005, 0.054) | 1.80E-02 | 0.01 (-0.034, 0.054) | 6.50E-01 | 0.025 (0.0035, 0.046) | 2.20E-02 | 0.0065 (0.00022, 0.013) | 4.30E-02 | 0.0048 (-0.039, 0.049) | 6.60E-01 | 0.0064 (0.00034, 0.012) | 3.80E-02 |
| Fasting proinsulin (pmol/L) | 0.015 (-0.05, 0.081) | 6.50E-01 | 0.084 (-0.034, 0.2) | 1.60E-01 | 0.032 (-0.026, 0.089) | 2.80E-01 | 0.0034 (-0.011, 0.018) | 6.50E-01 | 0.04 (-0.078, 0.16) | 2.30E-01 | 0.0051 (-0.0091, 0.019) | 4.80E-01 |
| 2h glucose (mmol/L) | -0.061 (-0.23, 0.1) | 4.70E-01 | -0.08 (-0.38, 0.22) | 6.00E-01 | -0.066 (-0.21, 0.079) | 3.70E-01 | -0.014 (-0.051, 0.024) | 4.80E-01 | -0.038 (-0.34, 0.26) | 6.10E-01 | -0.015 (-0.051, 0.021) | 4.10E-01 |
| HOMA-IR ((mU/L)*(mmol/L)) | 0.027 (-0.0087, 0.062) | 1.40E-01 | 0.041 (-0.027, 0.11) | 2.40E-01 | 0.03 (-0.0016, 0.061) | 6.30E-02 | 0.0059 (-0.0025, 0.014) | 1.70E-01 | 0.019 (-0.048, 0.087) | 3.00E-01 | 0.0066 (-0.0016, 0.015) | 1.10E-01 |
| HOMA-B ((mU/L)/(mmol/L)) | 0.0027 (-0.027, 0.032) | 8.60E-01 | 0.022 (-0.032, 0.077) | 4.30E-01 | 0.007 (-0.019, 0.033) | 5.90E-01 | 0.00059 (-0.0059, 0.0071) | 8.60E-01 | 0.011 (-0.044, 0.065) | 4.50E-01 | 0.0011 (-0.0052, 0.0074) | 7.30E-01 |
| AUCins (mU*min/L) | -0.025 (-0.22, 0.17) | 8.00E-01 | 0.033 (-0.33, 0.4) | 8.60E-01 | -0.012 (-0.18, 0.16) | 8.90E-01 | -0.0056 (-0.049, 0.038) | 8.00E-01 | 0.016 (-0.35, 0.38) | 8.60E-01 | -0.0043 (-0.046, 0.038) | 8.40E-01 |
| AUCins/AUCgluc (mU/mmol) | -0.022 (-0.22, 0.17) | 8.30E-01 | 0.031 (-0.33, 0.39) | 8.70E-01 | -0.01 (-0.18, 0.16) | 9.10E-01 | -0.0048 (-0.048, 0.038) | 8.30E-01 | 0.015 (-0.35, 0.38) | 8.70E-01 | -0.0037 (-0.046, 0.038) | 8.60E-01 |
| Incre30 (mU/L) | -0.00057 (- | 1.00E+00 | -0.0076 | 9.70E-01 | -0.0021 (-0.17, 0.16) | 9.80E-01 | -0.00013 (-0.042, 0.041) | 1.00E+00 | -0.0036 (-0.36, 0.35) | 9.70E-01 | -0.00033 (-0.041, 0.04) | 9.90E-01 |

| | | | | | | | | | | | | |
|---|---|---|---|---|---|---|---|---|---|---|---|---|
| | 0.19, 0.19) | | (-0.36, 0.35) | | | | | | | | | |
| Ins30adjBMI | -0.01 (-0.21, 0.18) | 9.20E-01 | 0.11 (-0.25, 0.46) | 5.50E-01 | 0.017 (-0.15, 0.19) | 8.40E-01 | -0.0023 (-0.045, 0.041) | 9.20E-01 | 0.052 (-0.3, 0.41) | 5.60E-01 | 0.00083 (-0.041, 0.043) | 9.70E-01 |
| ISI (mg/dL) | 0.11 (-0.1, 0.33) | 3.00E-01 | -0.28 (-0.64, 0.086) | 1.30E-01 | 0.011 (-0.18, 0.2) | 9.10E-01 | 0.025 (-0.024, 0.075) | 3.20E-01 | -0.13 (-0.49, 0.23) | 2.10E-01 | 0.017 (-0.032, 0.065) | 5.00E-01 |
| CIRadjBMI | -0.024 (-0.22, 0.17) | 8.10E-01 | 0.055 (-0.3, 0.41) | 7.60E-01 | -0.0055 (-0.18, 0.17) | 9.50E-01 | -0.0052 (-0.048, 0.038) | 8.10E-01 | 0.026 (-0.33, 0.38) | 7.60E-01 | -0.0034 (-0.045, 0.039) | 8.80E-01 |
| DI | 0.11 (-0.066, 0.3) | 2.10E-01 | -0.05 (-0.39, 0.29) | 7.70E-01 | 0.078 (-0.082, 0.24) | 3.40E-01 | 0.025 (-0.017, 0.067) | 2.40E-01 | -0.024 (-0.36, 0.31) | 7.70E-01 | 0.022 (-0.018, 0.063) | 2.80E-01 |
| **Obesity traits** | | | | | | | | | | | | |
| BMI (SD) | -0.031 (-0.048, -0.015) | 2.00E-04 | -0.017 (-0.043, 0.0092) | 2.00E-01 | -0.027 (-0.041, -0.013) | 1.30E-04 | -0.0069 (-0.012, -0.0019) | 6.70E-03 | -0.008 (-0.034, 0.018) | 2.70E-01 | -0.0071 (-0.012, -0.0023) | 3.40E-03 |
| WHR (SD) | 0.011 (-0.0055, 0.028) | 1.90E-01 | 0.056 (0.025, 0.088) | 4.90E-04 | 0.021 (0.0062, 0.036) | 5.40E-03 | 0.0025 (-0.0014, 0.0063) | 2.10E-01 | 0.027 (-0.0049, 0.058) | 5.80E-02 | 0.0029 (-0.00091, 0.0067) | 1.30E-01 |
| WHRadjBMI (SD) | 0.03 (0.013, 0.047) | 6.50E-04 | 0.075 (0.04, 0.11) | 2.40E-05 | 0.039 (0.023, 0.054) | 8.20E-07 | 0.0066 (0.0016, 0.012) | 9.80E-03 | 0.036 (0.00086, 0.07) | 4.60E-02 | 0.0072 (0.0022, 0.012) | 4.50E-03 |
| HDL (SD) | -0.06 (-0.1, -0.016) | 8.20E-03 | 0.023 (-0.033, 0.078) | 4.20E-01 | -0.028 (-0.062, 0.0071) | 1.20E-01 | -0.013 (-0.025, -0.0014) | 2.80E-02 | 0.011 (-0.045, 0.066) | 4.50E-01 | -0.0096 (-0.021, 0.0013) | 8.30E-02 |
| LDL (SD) | -0.058 (-0.11, -0.0097) | 1.90E-02 | -0.44 (-0.58, -0.3) | 7.60E-10 | -0.098 (-0.14, -0.053) | 2.30E-05 | -0.013 (-0.025, -4e-04) | 4.30E-02 | -0.21 (-0.35, -0.068) | 3.40E-02 | -0.014 (-0.026, -0.0012) | 3.10E-02 |
| TC (SD) | -0.084 (-0.13, -0.036) | 6.80E-04 | -0.51 (-0.67, -0.35) | 2.20E-10 | -0.12 (-0.17, -0.074) | 3.30E-07 | -0.019 (-0.033, -0.0044) | 9.90E-03 | -0.24 (-0.4, -0.085) | 3.30E-02 | -0.019 (-0.033, -0.0054) | 6.80E-03 |
| TG (SD) | 0.02 (-0.023, 0.062) | 3.60E-01 | -0.44 (-0.58, -0.31) | 3.50E-10 | -0.02 (-0.06, 0.021) | 3.40E-01 | 0.0043 (-0.0052, 0.014) | 3.80E-01 | -0.21 (-0.35, -0.072) | 3.30E-02 | 0.0038 (-0.0057, 0.013) | 4.40E-01 |

Effect was estimated by Wald’s method and inverse variance weighted (IVW) method for single variant, combined variants, respectively.

T2D: type 2 diabetes; HOMA-IR: The homeostatic model assessment (HOMA) insulin resistance; HOMA-B: HOMA beta cell function; AUCins: area under the curve (AUC) of insulin levels during oral glucose tolerance test (OGTT); AUCins/AUCgluc: ratio of AUC insulin and AUC glucose; Incre30: incremental insulin at 30 min;

Ins30adjBMI: Insulin response to glucose during the first 30 min adjusted for BMI; ISI: insulin sensitivity index; CIRadjBMI: Corrected Insulin Response adjusted for ISI; DI: disposition index; BMI: body mass index; WHR: waist-hip ratio; WHRadjBMI: WHR adjusted for BMI; HDL: high-density lipoprotein cholesterol; LDL: low-density lipoprotein cholesterol; TC: total cholesterol; TG: triglycerides; OR: odds ratio; CI: confidence interval; SD: standard deviation.

**Supplemental Table 5.** MR estimates following outlier removal

| | Initial Estimate | | | | | | After outlier removal | | | | | |
|---|---|---|---|---|---|---|---|---|---|---|---|---|
| **Trait** | **# SNPs** | **F** | **IVW** | | **Pleiotropy test** | | **# SNPs** | **F** | **IVW** | | **Pleiotropy test** | |
| | | | **Effect (95% CI)** | ***p*** | **MR-PRESSO global test p** | **modified Q' *p*** | | | **Effect (95% CI)** | ***p*** | **MR-PRESSO global test p** | **modified Q' *p*** |
| T2D (logOR) | 318 | 69 | 1.14 (1.05, 1.22) | 9.60E-04 | 1.50E-03 | 7.34E-04 | 315 | 69 | 1.1 (1, 1.2) | 1.67E-03 | 3.11E-01 | 3.26E-01 |
| Fasting insulin (pmol/L) | 10 | 37 | 2.36 (0.44, 12.18) | 3.15E-01 | 2.00E-04 | 3.14E-04 | 9 | 36 | 7.7 (1.3, 46) | 2.40E-02 | 3.18E-01 | 2.82E-01 |
| BMI (SD) | 1843 | 60 | 2.27 (1.95, 2.66) | 4.31E-25 | <2E-04 | 1.55E-04 | 1839 | 60 | 2.3 (2, 2.7) | 1.41E-25 | 2.50E-03 | 3.89E-03 |
| HDL (SD) | 228 | 123 | 0.88 (0.76, 1.02) | 7.78E-02 | <2E-04 | 2.18E-05 | 226 | 123 | 0.88 (0.76, 1) | 8.78E-02 | 2.00E-03 | 1.26E-03 |
| LDL (SD) | 193 | 149 | 0.93 (0.81 1.07) | 2.95E-01 | <2E-04 | 1.11E-07 | 192 | 148 | 0.96 (0.84, 1.1) | 5.47E-01 | <5.00E-04 | 7.29E-06 |
| TC (SD) | 242 | 112 | 0.92 (0.80, 1.06) | 2.38E-01 | <2E-04 | 4.23E-07 | 239 | 110 | 0.94 (0.82, 1.1) | 4.18E-01 | 1.00E-03 | 6.33E-04 |
| TG (SD) | 151 | 120 | 1.48 (1.22 1.77) | 3.74E-05 | <2E-04 | 7.04E-06 | 149 | 119 | 1.6 (1.3, 1.9) | 5.10E-07 | 7.00E-03 | 7.83E-03 |

IVW: inverse variance weighted; F: F statistics for the strength of correlation between instrument and exposure; MR-PRESSO: MR pleiotropy residual sum and outlier; Q’: Q’ statistics with modified second order weights; T2D: type 2 diabetes; BMI: body mass index; HDL: high-density lipoprotein cholesterol; LDL: low-density lipoprotein cholesterol; TC: total cholesterol; TG: triglycerides; OR: odds ratio; CI: confidence interval; SD: standard deviation.

**Supplemental Table 6**. Characteristics of the associations of PNPLA3 rs738409, NCAN rs2228603, and TM6SF2 rs58542926 with NAFLD in UKBB samples

| Gene | SNP | logOR | SE | P value |
|---|---|---|---|---|
| PNPLA3 | rs738409 | 0.33 | 0.053 | 2.09E-10 |
| NCAN | rs2228603 | 0.32 | 0.081 | 6.83E-05 |
| TM6SF2 | rs58542926 | 0.39 | 0.082 | 2.69E-06 |

OR: Odds ratio; SE: standard error.
